# Organisms associated with siphonophores revealed by short-read whole-genome sequencing

**DOI:** 10.64898/2026.10.29.762746

**Authors:** Casey W. Dunn, Namrata Ahuja, Dalila Destanović, Maciej K. Mańko, Martina Dal Bello, Samuel H. Church

**Author notes:** Corresponding author: Casey W. Dunn.

## Abstract

Siphonophores are associated with prey, parasites, microbes, and other organisms in the open ocean, but little is known about these siphonophore cobionts. To learn more about them, we investigated molecular bycatch: their DNA captured during siphonophore genome sequencing. We reanalysed 213 specimens of *Physalia, Nanomia, Bargmannia* and other siphonophores sequenced in four previously published studies. We recovered 104 bacterial genomes forming 32 genome clusters, including recurrent Metamycoplasmataceae and diverse proteobacteria, groups also found with other cnidarians. Bacterial communities differed among *Physalia* species sampled in the southwestern Pacific. We identified non-siphonophore eukaryotes from ribosomal markers in 38 specimens, including fish and copepods consistent with prey, parasitic flatworms, and *Perkinsela*-like relatives of amoeba endosymbionts. Alfonsino (*Beryx*) sequences in one Hawaiian *P. utriculus* specimen are consistent with capture of pelagic young of deepwater fishes. Molecular bycatch reveals the diversity of siphonophore associates from specimens already collected for genome research.

## Introduction

Siphonophores (Cnidaria: Hydrozoa) are abundant gelatinous predators in the ocean. They eat fish, crustaceans and other zooplankton, with marked prey differences among siphonophore groups [1, 2]. Siphonophores also host organisms on their surfaces and within their tissues. These associates, often termed cobionts [3], connect siphonophores to ocean food webs through feeding and other interactions.

Most studies of cnidarian microbiomes have focused on benthic animals. The Cnidarian Microbiome Database, for example, is dominated by surveys of stony corals, octocorals and sea anemones [4]. Microbial surveys of jellyfish and genomes recovered from octocorals provide useful comparisons for siphonophores [5–7], whose microbial associations remain poorly characterized. Studying these associates can also reveal organisms that are difficult to sample directly. Predators have, for example, been used to study poorly sampled midwater prey communities [8].

DNA sequenced for an animal genome often includes DNA from cobionts. Previous researchers have recovered microbial communities from mosquito genome datasets, *Wolbachia* genomes from biodiversity sequencing, and food plants, pathogens and symbionts from insect genome skims [9–11]. Parasites and other associates have also been separated from long-read genome data [3]. Holmes and Davis Rabosky called the incidental recovery of such natural history information from RADseq data “bycatch” [12]. This molecular bycatch can yield markers for identification and genomes for comparing the evolution and biology of associates.

Which organisms recur with siphonophores, and how do cobiont distributions vary among hosts and places? To address these questions, we reanalysed specimens sequenced in four studies on *Physalia* population genomics, *Nanomia* genome structure, *Bargmannia* diversity and siphonophore genome skimming [13–16]. The samples span surface-dwelling and water-column siphonophores and several ocean regions. We combined marker-based identifications with reconstructed bacterial genomes to identify these associates and compare their relationships and distributions.

## Results

### Human and laboratory-associated sequences

Human-compatible reads and bacteria associated with skin or laboratory reagents occurred in the sequencing data at a very low frequency in a subset of samples. The human screen retained 4,563 of 213,000,000 screened read pairs; the highest fraction was in *N. bijuga* NA31 (0.1058%). Skin-associated *Cutibacterium* and *Lawsonella* and reagent-associated bacteria were present, together with a *Serratia* clone and SSUs identical to *Caenorhabditis elegans*. Both libraries with these SSUs, TMAG-K5632 and TMAG-K5633, were scored as moderate quality by Church and colleagues [13]. We excluded four libraries from biological analyses: *Nanomia* sp. 1 NA29, *N. bijuga* NA31, *B. stenotes* D666-D10 and the formalin-preserved *B. elongata* T676-SS3. They met thresholds of at least 50% *C. acnes* breadth or 0.05% human-compatible read pairs, or exceeded the SSU background calibration. We set the thresholds after inspecting gaps in the distributions (Fig H in S1 Appendix). In the remaining 209 specimens, we set these putative contaminant taxa aside from biological summaries while retaining their flagged detections in the supporting tables. All *Physalia* libraries, including the ten scored as moderate quality by Church and colleagues, were retained under these criteria.

### Genome sequencing reveals a broad assemblage of associated organisms

DNA sequenced from siphonophores revealed bacteria, viruses and diverse eukaryotes (Fig 1). The recovered bacteria included groups known from other cnidarians, while eukaryotic markers identified likely prey, parasites and organisms carried by other associates (Table 1). Sampling spanned surface-dwelling and water-column siphonophores and one epibenthic specimen, with most specimens from *Physalia* and *Nanomia* (Figs A and B in S1 Appendix).

**Table 1.** Evidence for associated organisms across 209 retained specimens.

| Evidence set | Targets | Detections | Specimens | Assembly support |
|---|---|---|---|---|
| Bacterial genomes | 166 | 507 | 131 | 294 |
| Viral sequence groups | 256 | 478 | 119 | 297 |
| Other eukaryotic groups | 15 | 57 | 38 | 47 |

Distinct targets are bacterial genomes, viral sequence groups (viral operational taxonomic units, vOTUs), or broad eukaryotic groups identified from small-subunit ribosomal RNA (SSU) markers. A detection is one target in one specimen; the last column counts detections with assembly support from the same specimen. For eukaryotes, 40 detections have reads + assembly; assembly support also includes assembled SSUs below the read threshold. Contaminant taxa are set aside. These categories differ in taxonomic resolution.

**Fig 1.**
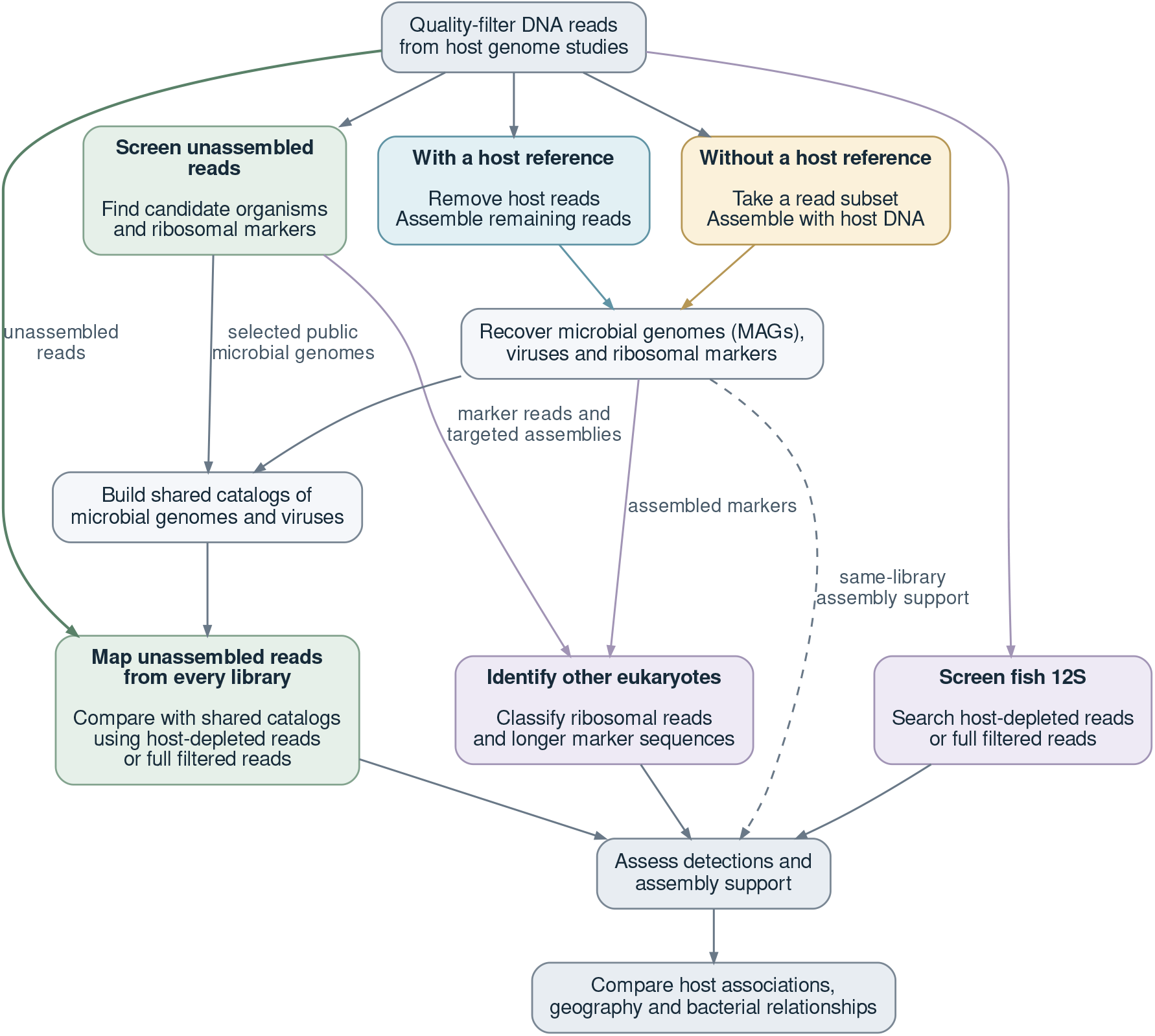
Identifying associates in DNA from 213 siphonophore specimens: 151 *Physalia*, 22 *Nanomia* and 40 other siphonophores. We screened unassembled reads and reconstructed metagenome-assembled genomes (MAGs). Reads matching siphonophore references were removed from *Physalia, Nanomia septata* and *Halistemma rubrum* libraries. We combined recovered sequences into shared catalogs and added selected public genomes to the bacterial catalog. We compared reads from every library with these catalogs and identified eukaryotes separately through ribosomal markers. A fish 12S screen searched host-depleted reads or full filtered reads from every library.

### Bacterial associates include diverse proteobacteria and recurrent Metamycoplasmataceae

Bacterial DNA was detected in 131 specimens, with matches to 166 genomes in our comparison catalog (Fig 2; Table 1). Proteobacteria (Pseudomonadota) included Rhodobacteraceae, *Pseudoalteromonas* and *Vibrio*. Other recovered groups included Bacteroidota, Actinomycetota and Cyanobacteriota.

**Fig 2.**
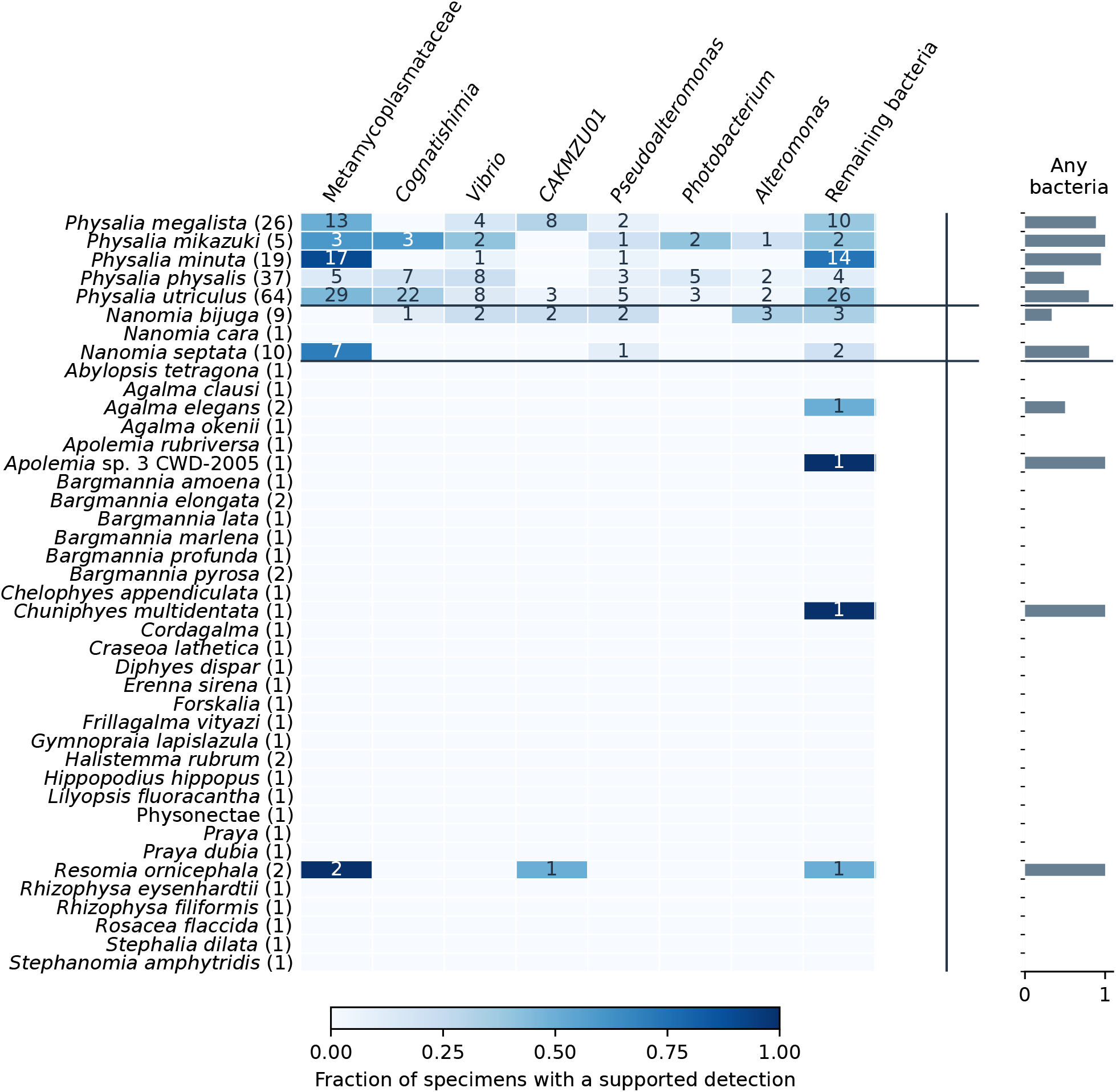
Proteobacteria and Metamycoplasmataceae recur across siphonophores. The seven displayed bacterial lineages are the most frequently detected identifiable groups, with Metamycoplasmataceae grouped together; other bacteria are pooled in a separate column. CAKMZU01 is an unnamed gammaproteobacterial lineage in GTDB. Colours show the fraction of specimens for each host identification with at least one accepted detection in the indicated microbial group. Each specimen contributes at most once to a group, even if it matches several genomes. Denominators include all retained libraries for that host, including those with no accepted microbial detection; contaminant taxa are set aside. Numbers in cells give positive specimens; host sample sizes appear in parentheses. The adjacent bars show the fraction with any bacterial detection.

Metamycoplasmataceae, a group of small, wall-less bacteria, were especially recurrent. We detected 6 genome clusters in 76 specimens. Counting each genome once per specimen gave 138 detections. Assembly of the original 213 libraries recovered 104 bacterial metagenome-assembled genomes (MAGs), forming 32 genome clusters; 55 met our near-complete criteria. A further genome belonged to the GTDB genus *Tyloplasma* (Hepatoplasmataceae), outside the Metamycoplasmataceae compared below (S1 Appendix).

### Vibrio occurs in *Physalia* and Nanomia

Read mapping detected *Vibrio* in 25 specimens: 23 *Physalia* and two *Nanomia bijuga* (NA33 and NA37). Two of these *Physalia* specimens yielded *Vibrio* MAGs: one assigned to *V. caribbeanicus* and the other to *V. alginolyticus*.

In some specimens, reads mapped to multiple related *Vibrio* reference genomes. Read mapping therefore did not resolve the identities or number of strains present (S1 Appendix).

### Siphonophore Metamycoplasmataceae have close relatives in other marine animals

The detected Metamycoplasmataceae occurred in *Physalia, Nanomia* and *Resomia* (Fig E in S1 Appendix). The seven reconstructed genome representatives belonged to a strongly supported group in the phylogeny of conserved bacterial markers (Fig 3; SH-aLRT 99.9%, ultrafast bootstrap 100%).

**Fig 3.**
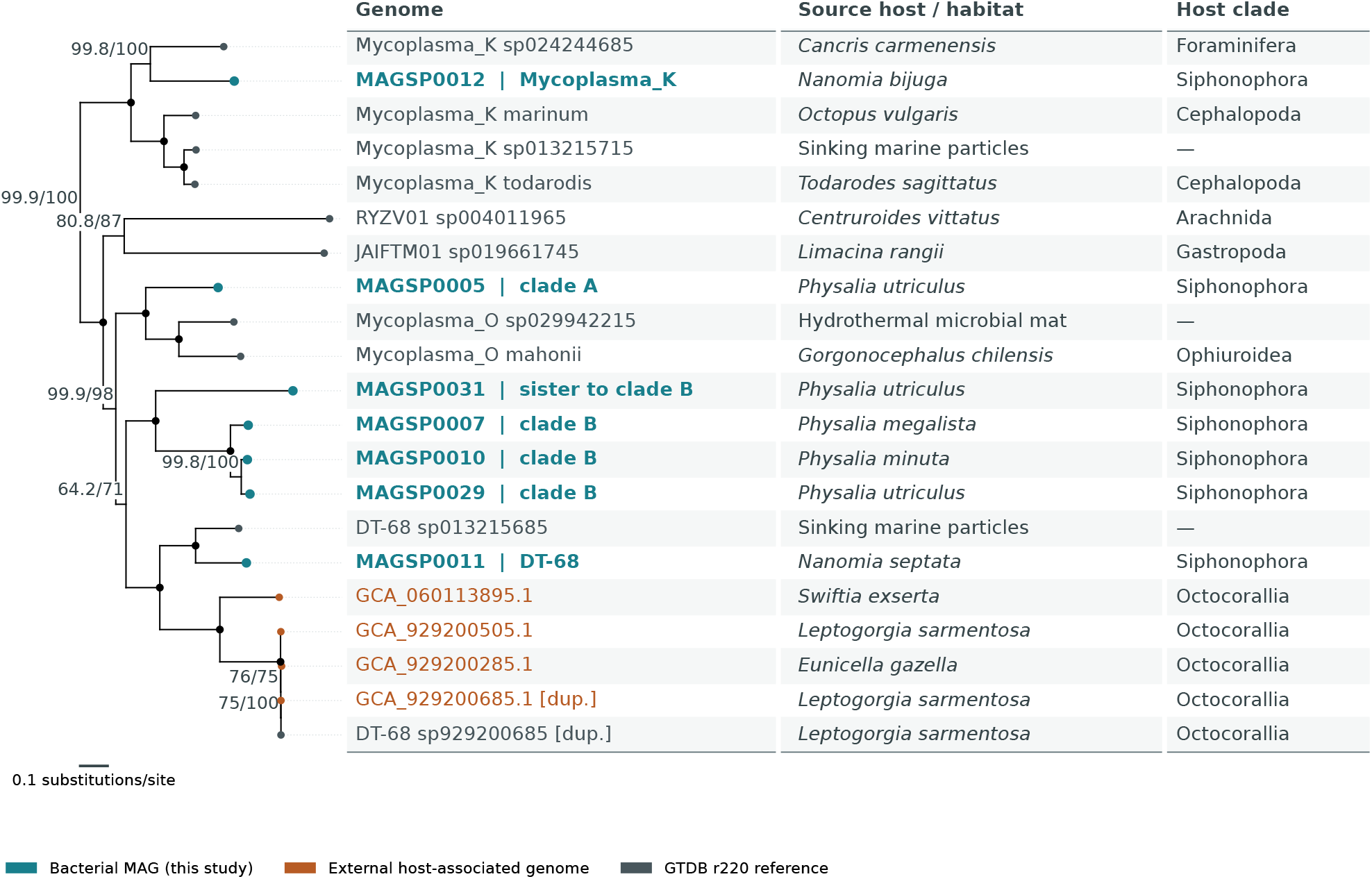
Siphonophore-associated Metamycoplasmataceae group with bacteria from diverse hosts and environmental samples. We inferred the tree with IQ-TREE 3 from 120 conserved bacterial marker proteins (bac120). The portion shown contains all seven siphonophore-associated Metamycoplasmataceae genome representatives and their closest sampled relatives. Branch lengths come from the full tree; node labels give SH-aLRT / ultrafast bootstrap support percentages; black internal dots indicate 100% for both measures. Tip colours distinguish genomes reconstructed here, genomes from other studies and GTDB reference genomes. The clade comprises 21 tips representing 20 distinct assemblies, including one assembly present in both external and GTDB inputs (marked [dup.]). The Source host / habitat column identifies the sampled organism or environment, and Host clade gives the corresponding host group. Dashes indicate environmental samples without an identified host. Accession-specific evidence from archive records and source publications is provided in the source-host table.

Clade A comprised MAGSP0005 and grouped with *Mycoplasma_O*, including a bacterium recovered from a basket star [17]. Clade B comprised MAGSP0007, MAGSP0010 and MAGSP0029; MAGSP0031 was a divergent sister lineage. The DT-68 genome, MAGSP0011, grouped with Metamycoplasmataceae genomes recovered from the octocorals *Eunicella gazella* and *Leptogorgia sarmentosa*, among others [7]. MAGSP0012 grouped with *Mycoplasma_K*, whose sampled relatives included bacteria from squid and octopus [18]. This genome was recovered from a single specimen, NA31, and could have arrived with prey. We retained it only for phylogenetic comparison after excluding NA31 from biological analyses. The clade-B/DT-68 sister relationship was weakly supported (SH-aLRT 64.2%, ultrafast bootstrap 71%).

The lineages also differed in their hosts. Clades A and B occurred only in *Physalia* in the retained data. DT-68 occurred in *N. septata* and *R. ornicephala* (Fig E in S1 Appendix).

Related bacteria also occurred in earlier cnidarian microbiome surveys. MAGs in five of the seven clusters contained assembled 16S ribosomal RNA genes. These matched sequences in 98 samples from 23 publications in the Cnidarian Microbiome Database, including cnidarian, sponge and seawater samples [4]. Amplicon sequence variants (ASVs) at least 99.5% identical to these assembled bacterial 16S ribosomal RNA genes accounted for at most 0.38% of a sample’s reads. More divergent relatives were sometimes abundant: variants 97.6% identical to MAGSP0005 made up most reads in low-depth libraries from laboratory-reared *Acropora loripes* and *Acropora sarmentosa* recruits in one study [19]. Independent datasets from the jellyfish *Cassiopea* yielded no match to the available MAG 16S sequences at the 90% reporting threshold (S1 Appendix).

### Bacterial communities differ among geographic regions and hosts

Three Metamycoplasmataceae genome clusters were frequent in southwestern Pacific and Indian Ocean *Physalia*, but rare in the central Pacific: MAGSP0005 in clade A, and MAGSP0007 and MAGSP0010 in clade B. All three varied significantly among regions after accounting for sequencing effort, with permutations within flowcells (S1 Appendix).

The southwestern-versus-central Pacific contrast persisted when the comparison was restricted to recorded tentacle samples (Fig D in S1 Appendix). Host identifications and collection histories still differed among regions.

The broader bacterial community also varied geographically. Of 122 *Physalia* specimens with recorded sampling regions and libraries each sequenced on a single flowcell, 88 had bacterial detections and entered the community comparison. Region accounted for 19.8% of the variation in which genomes were detected, after accounting for the number of input reads (Jaccard PERMANOVA, *P* ≤ 0.0002). The regional association persisted under both alternative detection thresholds (Table A in S1 Appendix). No testable genome showed significant concentration on particular flowcells after host and region were taken into account (S1 Appendix).

Within southwestern Pacific tentacle samples, bacterial communities differed among *P. megalista, P. minuta* and *P. utriculus* with input reads and flowcell as covariates and permutations within flowcells (30 specimens; *R*^2^ = 13.3%, *P* ≤ 0.0002). This difference persisted at both alternative detection thresholds (S1 Appendix). Within *P. utriculus* tentacles, no regional difference was detected among the central, northwestern and southwestern Pacific at the primary threshold (31 specimens; *R*^2^ = 6.1%, *P* = 0.2916). This regional comparison was significant only at the 5% breadth threshold (S1 Appendix).

### Uneven sampling limits comparisons among tissues and depths

Most samples were *Physalia* tentacles or siphosomes from other siphonophores. The few alternative tissues were unevenly distributed among hosts, regions and flowcells, preventing a general test of tissue effects. Fish 12S matches occurred in 2 of 3 zooid samples and 6 of 142 tentacle samples. These sparse observations suggest that the tissue sampled influences prey recovery. The two positive zooid samples came from the same northwestern Hawaiian locality and shared sequencing lanes.

Depth and geography were also confounded (Figs C and G in S1 Appendix). Within *N. septata*, the Metamycoplasmataceae bacterial lineage DT-68 was detected in all seven California specimens but none of the three from Washington. Because the Washington specimens were collected near the surface (0–20 m) and the California specimens at 217–662 m, this difference cannot be attributed separately to geography or depth.

### Viral sequences include bacteriophages and endogenous candidates

Most recovered viral groups belonged to Uroviricota, which contains tailed bacteriophages. Other groups included Cressdnaviricota and larger DNA viruses. We detected 256 associate viral groups in 119 specimens (Table 1). Sequences that could derive from host genomes, including possible integrated copies of RNA viruses, were recorded separately as endogenous candidates.

### Eukaryotic associates include prey and parasite lineages

Eukaryotic markers identified likely prey and parasites alongside amoebae, kinetoplastids, fungi, dinoflagellates, algae and ciliates. We detected 15 broad groups in 38 specimens (Fig 4; Table 1). Of the 57 detections, 47 were assembly-supported, including 40 with both reads and assembly.

**Fig 4.**
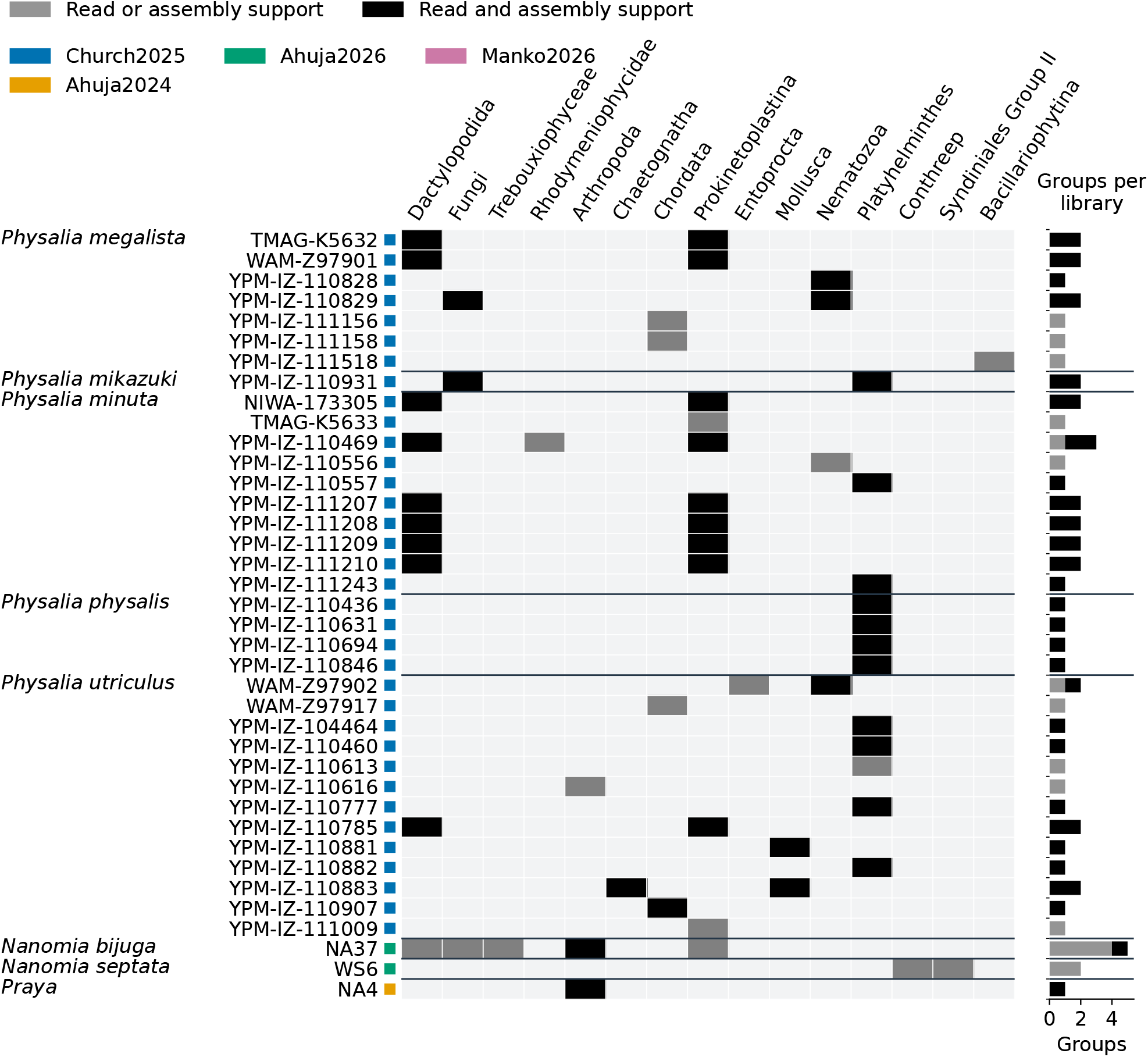
Eukaryotic associates include animals, protists and fungi. Rows show the 38 specimens with at least one accepted detection, grouped by host identification. Black cells combine read and assembly support; medium-grey cells have one of those forms of support. Light-grey cells do not meet the reported presence criterion. Host names and specimen identifiers appear at left; each host name aligns with the first specimen in its group. Coloured squares identify source studies. Columns show broad eukaryotic groups, with animals grouped by phylum. Conthreep is a group of ciliates in the SILVA classification. Bars at right count detected groups per specimen, separated by the type of evidence. Each group is counted once per specimen, giving 57 detections; overlapping classifications are counted at the more specific level. The display includes retained specimens and sets contaminant taxa aside. Full identities and ecological annotations are in the supporting evidence table.

*Physalia, Erenna* and *Rhizophysa* are known to feed on fish [1, 20, 21]. Crustaceans are important prey for many other siphonophores [2]. Our data included nine detections of likely prey across eight specimens. Fish 18S ribosomal sequences occurred in two *Physalia utriculus* and two *P. megalista*. An assembled fish 18S sequence in one of the Hawaiian *P. utriculus* specimens was corroborated by the mitochondrial 12S evidence described below (S1 Appendix).

Assembled arthropod ribosomal sequences from one *Praya* sp. and one *Nanomia bijuga* matched the copepods *Metridia* and *Euchaeta*, respectively. Bivalve sequences occurred in two *P. utriculus* specimens, with chaetognath sequences in one of the same specimens.

A dedicated fish 12S screen across all 213 libraries recovered 32 long, close fish-matching read pairs in 8 *Physalia* specimens. Reference affinities included anchovies (*Engraulis*), alfonsinos (*Beryx*) and carangids, with near-tied matches limiting species identification. The *Beryx* matches occurred in the Hawaiian *P. utriculus* with the assembled fish 18S sequence. No other siphonophore group yielded qualifying fish matches, although known fish-feeding groups were sparsely sampled (one *Erenna* and two *Rhizophysa*; S1 Appendix).

Parasitic flatworms occurred in 11 *Physalia* specimens. This evidence comprised 12 assembled ribosomal sequences. Trematodes (flukes) occurred in eight specimens and cestodes (tapeworms) in four; one *P. physalis* contained both. Trematode sequences occurred in *P. utriculus, P. physalis, P. minuta* and *P. mikazuki*, with closest references including *Dinurus, Preptetos* and *Prodistomum*. Cestode sequences occurred in two *P. utriculus*, one *P. physalis* and one *P. minuta*; the last closely matched *Echinophallus wageneri*.

Additional markers allowed direct comparison with trematodes collected by Waki and colleagues [22]. In *P. utriculus* from Hawai’i and *P. mikazuki* from Japan, these ITS2-containing and mitochondrial COI sequences closely matched *Dinurus tornatus*. An ITS2-containing region from *P. physalis* in Florida was closest to *D. barbatus*. In two *P. physalis* from the northeastern United States, 28S sequences were closer to Waki’s *Prodistomum* Type 4 than to *P. orientale*, whereas COI sequences were more divergent (S1 Appendix).

Eleven detections involved relatives of *Perkinsela*, a kinetoplastid endosymbiont of *Paramoeba* amoebae. Nine of these ribosomal detections occurred in the same specimens as accepted dactylopodid amoeba sequences, consistent with this nested association.

Four related genomic bins also matched *Perkinsela*. These bins were initially classified as archaeal, but protein comparisons showed they were eukaryotic. We therefore excluded them from prokaryotic counts and retained them for eukaryotic study (S1 Appendix).

### Assembly recovered bacteria absent from reference databases

Assembly recovered 24 catalog representatives absent from the reference databases used by sylph, including the focal Metamycoplasmataceae. Bracken and phyloFlash had detected broader mycoplasma signals in those specimens. General assembly also recovered additional eukaryotic markers, including a *Perkinsela*-like sequence missed by the phyloFlash assembly and read-presence criteria (S1 Appendix).

### Host DNA removal changed some detections

In species where a reference genome was available for identifying host sequences, a large fraction of reads were removed before cobiont assembly: a median of 98.53% in *Physalia*, 99.90% in *N. septata* and 99.23% in *H. rubrum*. This predominance of host DNA reflects the original studies’ focus on host genome sequencing. A comparison of 10 libraries that held the bacterial and viral reference catalogs constant found that most bacterial detections were shared between host-depleted and full-read inputs. Full reads yielded more viral detections in that comparison, which included both associate and endogenous-candidate sequences (S1 Appendix).

## Discussion

### Siphonophores share bacterial groups with other cnidarians

Several prominent bacterial groups found here also occur with other marine cnidarians. Rhodobacteraceae, *Pseudoalteromonas* and *Vibrio* have been recovered from living *Aurelia* medusae, including cultures from their external surfaces [6]. Earlier targeted work detected *Tenacibaculum maritimum* DNA with the siphonophore *Muggiaea atlantica* [23]. Our survey extends these observations with bacterial genomes from multiple siphonophore lineages.

A cnidarian-associated bacterium can also have different effects under different conditions. Vibrios occur in the microbiota of *Aurelia* and *Cassiopea* [6, 24]. *In the coral Pocillopora damicornis, V. coralliilyticus* swims toward chemical cues in mucus, with stronger attraction to mucus from heat-stressed corals [25]. During infection it damages coral tissues and disrupts nutrient exchange with algal symbionts [26]. The *Vibrio* genomes recovered from *Physalia*, together with detections in *Nanomia*, raise questions about harmless colonization, opportunistic disease and carriage with prey in siphonophores.

Our findings extend the bacterial screening reported by Church and colleagues [13]. Their Kraken2 screen identified strong Vibrionales signals in three *Physalia* libraries that were excluded before defining the 151-library dataset reanalysed here. We detected *Vibrio* in 23 retained specimens and recovered genomes assigned to *V. caribbeanicus* and *V. alginolyticus* from two of them. *Vibrio* sequences were therefore present beyond the libraries flagged by the original study. These findings are qualitatively consistent with their screening, while assembly and genome mapping provide additional information about bacterial identities and distributions. Our detection frequency applies to the previously filtered cohort; a systematic comparison with their original bacterial assignments would be needed to quantify agreement between the analyses.

### Host identity and geography structure bacterial associations

Within *P. utriculus*, three Metamycoplasmataceae genome clusters, from two lineages, were frequent in southwestern Pacific and Indian Ocean specimens but rare in the central Pacific. Clades A and B occurred in *Physalia*, whereas DT-68 occurred in *N. septata* and *R. ornicephala*. Sampling the same hosts across regions and depths would help separate these influences on bacterial distributions.

Collection history may also contribute to the differences among hosts. Many *Physalia* were collected on beaches and preserved in ethanol, whereas many other specimens were collected alive and flash frozen. Bacteria could have grown in beached *Physalia* before preservation. We detected *Pseudoalteromonas, Alteromonas* and *Vibrio* in *Physalia*. These bacteria rapidly use jellyfish detritus [27], so their occurrence in these specimens could reflect associations during life, growth after death, or both. Incomplete preservation histories and the scarcity of comparable tissues limit our ability to separate these possibilities.

### Prey sequences connect molecular bycatch to feeding observations

Fish and copepod sequences agree with known differences in siphonophore diets. Fish are the principal prey of *Physalia*, whose recorded prey also include chaetognaths, or arrow worms [1]. Their recovery here corroborates those observations. Copepod-like sequences in *Praya* sp. and *Nanomia bijuga* fit the importance of crustaceans in other siphonophore diets [2].

Fish sequences also suggest prey whose adults occupy deeper habitats. Mitochondrial sequences from a Hawaiian *P. utriculus* matched alfonsinos (*Beryx*), which live near the seafloor in deep water as adults [28]. Their larvae have been collected primarily in the upper 50 m near Hancock Seamount [29]. These matches add alfonsinos to the potential prey of *Physalia* and are consistent with capture of their pelagic young.

The same specimen yielded an 18S sequence whose closest archived reference was *Centroberyx gerrardi*, a fish recorded from temperate southern Australian waters [30]. This conserved 18S sequence supports only a broad fish identification; it does not establish that *C. gerrardi* occurs in Hawai’i.

Published observations describe *Nomeus* swimming beneath *Physalia* and consuming its zooids and tentacles [31]. We did not identify *Nomeus* in our sequence data. Fish DNA can thus record feeding by the siphonophore or other encounters with fish.

Bivalve sequences in *P. utriculus* could reflect direct capture of planktonic life stages or ingestion within another animal’s gut. Such secondary predation likely explains crustaceans in historical *Physalia* gut samples [1] and has been considered for small prey signals in dietary metabarcoding [2]. That study also recovered a ctenophore from *Physalia* but could not distinguish direct capture from secondary predation. Molecular bycatch can therefore record more than one feeding step.

### Parasites connect siphonophores with other marine hosts

Fluke and tapeworm sequences extend the molecular documentation of siphonophore parasites. Larval flukes have been observed within siphonophore tissues [32], and gut-content metabarcoding recovered both groups [2]. Our assembled markers identify flukes in four named *Physalia* taxa and tapeworms in three.

The additional nuclear and mitochondrial markers connect these detections to parasite life cycles. Waki and colleagues matched *Dinurus tornatus* larvae in *Physalia* to adults in mahi-mahi, *Coryphaena hippurus*; their *Prodistomum orientale* comparisons also included adults from mahi-mahi and chub mackerel, *Scomber japonicus* [22]. Our Hawaiian *P. utriculus* and Japanese *P. mikazuki* contained sequences closely matching their *D. tornatus*. The Florida *D. barbatus*-like sequence and northeastern US *Prodistomum*-like sequences broaden the material available to trace transmission when fish eat gelatinous hosts, although the differing nuclear and mitochondrial affinities of the latter leave their identity unresolved.

Sampling for host genomes can miss conspicuous associates. Hyperiid amphipods attach to siphonophores and feed on their tissues [33], yet we found no hyperiid sequences. They could have been removed before host-tissue sampling. Other eukaryotes, including fungi, green algae, diatoms and ciliates, have less resolved roles: they may have been attached, ingested or collected alongside the siphonophores, or carried within ingested prey. Targeted sampling and direct observations would help distinguish these possibilities.

### Amoebae may carry their own endosymbionts

The repeated pairing of amoebae with *Perkinsela*-like sequences suggests that some eukaryotic associates live within others. *Perkinsela* lives inside *Paramoeba* amoebae [34]. Amoebae related to *Paramoeba* occurred in ten specimens, nine alongside *Perkinsela*-like sequences; two further specimens contained only the latter. Direct observation is needed to confirm the intracellular relationship here and locate the amoebae relative to the siphonophore. The related genomic bins offer further material for studying these organisms.

### Bacterial associations and the potential of molecular bycatch

Bacterial detections were unevenly distributed among siphonophores, and more than one-third of specimens yielded no accepted detection (Fig 2). Comparisons with other cnidarians provide context for these sparse detections. The Cnidarian Microbiome Database revealed less diverse microbiomes in sampled hydrozoans and jellyfish than in stony corals [4]. Our marker comparisons also identified relatives of siphonophore-associated Metamycoplasmataceae in other marine animals. Whether siphonophores support less diverse communities or lower bacterial loads remains unresolved: targeted amplicon surveys and host genome sequencing differ in their recovery of microbial diversity. Consistent tissue sampling, deeper microbial sequencing and quantitative measurements of bacterial abundance would help distinguish biological differences from incomplete detection.

Our analyses yielded new biological insight into siphonophore feeding, parasitism and bacterial associations from datasets collected to study the hosts themselves. Studies designed specifically for cobionts would use sampling suited to their questions, such as gut tissue for prey detection or surface swabs for bacteria on the animals’ exterior, and apply consistent sampling across specimens for microbiome comparisons. Such studies often use amplicon sequencing to reduce costs and sequence markers from selected groups more deeply. In our repurposed datasets, inconsistent tissue sampling and uneven sequencing depth for cobionts make it difficult to distinguish true absences from missed detections. Shotgun sequencing has complementary advantages: it can recover sequences from diverse organisms without specifying target groups in advance, providing genomic information beyond marker loci. Molecular bycatch remains an underused source of biological information. As genomic datasets accumulate, their bycatch will provide an increasingly valuable resource for studying cobionts and their interactions with hosts.

## Materials and Methods

### Ethics statement

This study consists of computational reanalysis of sequencing data from previously published studies.

### Sampling associated organisms through siphonophore genome sequencing

We reanalysed short-read DNA sequencing from four siphonophore genome studies [13–16], with one sequencing library per specimen. *Physalia* were typically collected from beaches and preserved in ethanol; many other siphonophores were collected alive by ROV or SCUBA and flash frozen.

Collection-to-preservation intervals and preservation details were not consistently recorded for individual specimens. The sample metadata table links each library to its source study, host identification, collection information and sequencing accessions. The 151 *Physalia* libraries exclude 12 removed by Church and colleagues: three with heavy Vibrionales signals, one with arthropod sequences, two with evidence of cross-sample contamination, and six replicates or duplicates (source-study QC report). We retained all ten libraries that they scored as moderate quality [13].

We identified candidate associates from reads, reconstructed genomes and checked their presence across specimens (Fig 1). We first subsampled libraries exceeding 200 million read pairs, then removed adapters and filtered reads with fastp [35]. We screened these reads with Kraken2, Bracken, sylph and phyloFlash [36–39].

For assembly, we removed reads matching siphonophore reference genomes from *Physalia, Nanomia septata* and *Halistemma rubrum*. Church and colleagues reported that 85–99% of reads mapped to a shared *P. physalis* reference across *Physalia* lineages [13]. Other *Nanomia* lineages mapped poorly to *N. septata*, so we retained host reads in those libraries and in hosts without reference genomes. Reference accessions and mapping criteria are given in S1 Appendix; the host-mapping table records the reads removed.

We attempted assembly of every library separately with MEGAHIT, using all remaining host-depleted reads or up to 25 million quality-filtered pairs when no host reference was used [40]. We pooled candidate microbial genomes across specimens with public reference genomes identified by sylph, then dereplicated them to form a shared catalog. Recovered viral sequences were pooled and dereplicated into a separate catalog. Reads from every library were mapped competitively to these catalogs, using all host-depleted reads where available and the full quality-filtered reads otherwise, subject to the initial 200-million-pair cap. Software settings and metadata procedures are detailed in S1 Appendix.

### Bacterial diversity and genome recovery

We reconstructed microbial genomes to identify associates. We binned contigs into metagenome-assembled genomes (MAGs) with MetaBAT2, assessed their quality with CheckM2 and classified them with GTDB-Tk [41–43]. We checked bins and contigs for misclassification and mixed origin using protein-domain searches and comparisons with host and other eukaryotic sequences. We reassessed quality after removing fragments identified as foreign (S1 Appendix). Retained genomes were at least 50% complete with less than 10% estimated contamination. We called genomes *near-complete* when these estimates were at least 90% completeness and below 5% contamination [44] (S1 Appendix). Genomes sharing at least 95% average nucleotide identity across at least half of either genome were grouped into species clusters using skani [45]. This genome-wide criterion follows the widely used 95% average nucleotide identity boundary [46]. Identifiers beginning MAGSP denote microbial genome clusters. We combined these reconstructed genomes with reference genomes identified by sylph, retaining one representative per cluster. The resulting comparison catalog contained 478 candidate genomes and 20 decoy genomes used to check for spurious mapping (S1 Appendix).

We competitively mapped reads from all libraries against the catalog using BWA-MEM [47], including libraries that yielded no MAG. After removing duplicates and poorly supported alignments, we measured the fraction of each genome covered by reads, termed *breadth*, with CoverM [48]. We accepted bacterial presence when at least 100 reads covered at least 10% of a genome; coverage of 1–10% was recorded as trace evidence. An accepted detection received additional assembly support if the same specimen yielded a reconstructed genome from that cluster, or contigs matching at least 10 kb of its catalog genome at 95% identity. These detections are labelled *assembly-supported* in the bacterial detection table. We compared assemblies with `minimap2 using -x asm5 -c –secondary=no` [49]. We checked recovery by mapping reads to genomes assembled from the same library and checked gross spurious mapping against distant decoy genomes. These controls did not calibrate confusion among close relatives or sensitivity at low abundance. One population can match several related references, so the number of genome matches is not a direct measure of species richness.

### Bacterial relationships

To place detected bacteria among their relatives, we inferred a phylogeny for the seven recovered Metamycoplasmataceae genome representatives and their relatives from other hosts and reference databases. The alignment used GTDB-Tk’s bac120 set of 120 conserved bacterial marker proteins [43]. We inferred the tree with IQ-TREE 3 and assessed branches with SH-aLRT and ultrafast bootstrap support [50–52]. We also compared available bacterial 16S ribosomal RNA genes with the Cnidarian Microbiome Database and independent *Cassiopea* datasets to identify related bacteria in earlier surveys [4, 24].

### Distributions among hosts and ocean regions

We described detections by host, collection depth and sampled tissue across the 209 retained specimens. Within *Nanomia*, we compared locality and depth with detections of a Metamycoplasmataceae bacterial lineage labelled DT-68 in the Genome Taxonomy Database (GTDB). Within *Physalia*, we repeated regional summaries using only recorded tentacle samples.

To assess sequencing-batch effects, we tested the 140 libraries that were sequenced on a single flowcell, including 122 *Physalia*. All retained libraries remained in descriptive summaries. We tested whether individual bacterial genomes were unusually concentrated on particular flowcells, using 10,000 permutations within host-by-region groups. Tests required sufficient replication; a false-discovery-adjusted *q* < 0.05 flagged possible contamination.

Tests of geographic variation were restricted to *Physalia*, with 11 named sampling regions spanning the Atlantic, Indian and Pacific oceans. For microbial genome clusters detected in at least ten eligible *Physalia* libraries, we tested whether detection varied among regions after accounting for the number of input read pairs. Because the planned logistic mixed models with a flowcell random effect fitted unreliably, we used the gain in logistic-model fit after adding region, assessed against 9,999 permutations of region labels within flowcells. To compare whole communities, we used Jaccard dissimilarity and PERMANOVA with the same within-flowcell permutation design. Input read count was entered before region, and libraries with no detected bacteria were excluded from this community test.

To separate host and regional patterns where sampling allowed, we made two comparisons restricted to tentacles: regions within *P. utriculus*, and *P. megalista, P. minuta* and *P. utriculus* within the southwestern Pacific. These targeted community tests included read count and flowcell as covariates, with permutations restricted within flowcells. Tissue and depth comparisons were descriptive where sampling did not permit their effects to be separated from host, region or batch (S1 Appendix). We checked sensitivity to detection thresholds, removal of host reads, input subsampling and the studies that contributed candidate genomes (S1 Appendix).

### Viral diversity

We identified viral candidates in the assemblies with geNomad and assessed their completeness with CheckV [53, 54]. We required at least one characteristic viral gene and either at least 5 kb of sequence or at least 50% estimated completeness. Sequences matching host genomes, or belonging to groups with plausible origins within host genomes, were recorded separately as *endogenous candidates* (S1 Appendix). We removed cellular flanks identified by CheckV and reapplied the inclusion criteria to the remaining viral regions. We grouped closely related sequences into viral operational taxonomic units (vOTUs), using 95% average nucleotide identity across at least 85% of the shorter sequence [55].

We mapped reads competitively against associate viruses and endogenous candidates together. Viral presence required coverage of at least 75% of a group’s representative sequence; 10–75% was a partial match. Recovery of a member of the same viral group from that specimen’s assembled sequences supplied additional support.

### Eukaryotic diversity, prey and parasites

We identified non-siphonophore eukaryotes using small-subunit ribosomal RNA gene (SSU; 18S in animals) sequences. Unassembled marker reads were compared against SILVA 138.1 supplemented with siphonophore SSU sequences assembled from the cohort [56]. When several references matched similarly well, the assignment was the narrowest taxonomic group they shared. We excluded cnidarian assignments and animal matches too broad to distinguish host from associate. This screen therefore cannot assess cnidarian prey. We accepted a detection supported by at least 37 read pairs at 97% identity or by a non-host assembled SSU. The read threshold was one pair above the maximum for selected control groups across 205 libraries (S1 Appendix). We retained this threshold across the cohort and flagged specimens exceeding the calibration background. Assembly-supported detections include both those with sufficient read evidence (“reads + assembly”) and those accepted from assembly alone.

To summarize eukaryotes without counting the same evidence repeatedly, we counted each broad group once per specimen. Animals were grouped by phylum; other eukaryotes used the first five levels of the SILVA classification. When both a group and one of its subgroups were detected, only the more specific assignment contributed to the total. The eukaryote evidence table retains all assignments. We checked assembled sequences against NCBI and used taxonomic identity and known biology to interpret likely prey, parasites and other associates.

We searched separately for fish mitochondrial 12S sequences, using BLASTN to compare candidate read pairs against MitoFish and mitochondrial RefSeq [57, 58]. We required long, close matches with no strong competing non-fish evidence. S1 Appendix describes the complementary search methods, criteria and controls.

For trematodes, we compared the existing assemblies of the eight positive specimens directly with sequences deposited by Waki and colleagues and references from their supplement [22, 58]. These comparisons used 28S ribosomal RNA, mitochondrial cytochrome c oxidase subunit I (COI), and regions containing the second internal transcribed spacer of ribosomal DNA (ITS2). The ITS2-containing sequences also include flanking ribosomal genes. We retained alignment lengths, identities and competing matches, and distinguished markers assembled on the same contig from those found separately in the same specimen (S1 Appendix).

### Human DNA and possible laboratory background

To assess human DNA, we screened one million raw read pairs per library, taking equal numbers from the beginning of each source file. After quality filtering, we aligned pairs to hg38 with Bowtie2, requiring concordant, full-length alignments of both mates at at least 99% identity and mapping quality 30. We tested the screen with simulated human and siphonophore read pairs, measuring recovery of human reads and misassignment of siphonophore reads. Candidate sequences were compared with the broader NCBI nt database (S1 Appendix). The resulting fractions describe these nonrandom subsets, which may differ from the complete libraries.

We handled possible contamination in two tiers. First, we excluded libraries with *C. acnes* breadth at least 50%, human-compatible read pairs at least 0.05% of screened pairs, or SSU background controls above the calibration range. These criteria excluded NA29, NA31, D666-D10 and T676-SS3. Second, we retained but flagged detections of skin- and reagent-associated groups and the laboratory-associated *Serratia* clone MAGSP0001 and *Caenorhabditis elegans* SSUs; both libraries with the latter sequences were scored as moderate quality by Church and colleagues [13]. These detections were set aside from all biological summaries and comparisons. The contaminant list uses published evidence for skin and reagent associations [59–61]; extraction blanks were unavailable. All *Physalia* libraries were retained.

## Abbreviations

ASV: amplicon sequence variant
GTDB: Genome Taxonomy Database
MAG: metagenome-assembled genome
PERMANOVA: permutational multivariate analysis of variance
SSU: small-subunit ribosomal RNA
vOTU: viral operational taxonomic unit.

## Use of generative artificial intelligence

Following the Generative AI Delegation Taxonomy (GAIDeT) [62], the authors used OpenAI Codex with Astra 6 for code generation and process automation in implementing analyses and orchestrating computational runs; data analysis and validation in assessing results; and text generation, proofreading and editing in drafting and revising the manuscript and S1 Appendix. Anthropic Claude Opus 5.5 was used for a technical review, corresponding to GAIDeT tasks of quality assessment, identification of limitations and recommendations. Outputs were checked through code tests, comparisons with source data and analysis results, primary-literature verification, visual inspection of figures and documents, and author review. The authors retain responsibility for the accuracy and originality of the analyses, interpretations, references and final manuscript.

## Funding Statement

Free access to OpenAI models was provided by OpenAI through the ChatGPT for Academic Researchers program. Free and reduced-rate access to Anthropic models was provided by Anthropic through the Claude Team plan for scientists program. OpenAI and Anthropic had no role in study design, data collection, analysis, interpretation, preparation of the manuscript, the decision to publish, or any other aspect of this work.

## Author contributions

Casey W. Dunn: conceptualization, methodology, formal analysis, writing—original draft, and writing—review and editing. Namrata Ahuja, Dalila Destanović, Maciej K. Mańko, Martina Dal Bello and Samuel H. Church: methodology (suggestions for analyses) and writing—review and editing.

## Data Availability

No new sequencing data were generated for this study. We reanalysed sequencing libraries from four published siphonophore studies [13–16]. The accompanying sample table links library and specimen identifiers to source studies, host identifications and recorded sequencing accessions. Supporting detection tables show which organisms were detected in each specimen and the evidence for each identification. Additional supporting files contain genome quality, statistical results and trematode-marker comparisons, with original analysis paths and checksums. Figure source tables and checksums accompany the manuscript assets. Analysis code, the executable workflow and figure-generation scripts are maintained at https://github.com/caseywdunn/siph-associates. An earlier version of the analysis code (v0.5.0) is archived at https://doi.org/10.5281/zenodo.23441461.

Submission of the reconstructed genomes, viral sequences, SSUs and marker contigs to NCBI is pending. The repository provides the preparation workflow, genome deposit decisions and provisional organism-name crosswalk, with submission instructions and current status. Sequence accession numbers and a DOI for the accompanying sequence archive will be added when available. Public sequencing accessions remain unverified for OA-190531-1 and OA-Bluebottle.

## Competing interests

The authors have declared that no competing interests exist.

## Supporting Information

**S1 Appendix. Supporting methods, results, figures and tables**. Sampling and sequence-analysis methods, detailed detection evidence, sensitivity analyses, collection context, bacterial relationships, and links to supporting data. (PDF)

## Notes

https://github.com/caseywdunn/siph-associates

