## Supporting information for "Organisms associated with siphonophores revealed by short-read whole-genome sequencing"

### S1 Appendix. Supporting information

#### S1 Supplementary methods

We identified associates by comparing reads and assembled sequences with reference genomes and ribosomal markers. Each library represents one siphonophore specimen. We count a detection once per organism or sequence group in a specimen, using the groupings and evidence thresholds specified below.

##### S1.1 Specimens, collection metadata and read preparation

We checked which specimens and sequencing libraries were represented across the four source studies [1–4] (source manifest and cohort manifest). The eight *Bargmannia* libraries are the specimens whose sequencing data are listed in Table S2 of Mańko and colleagues [4], including *B. marlena* and *B. pyrosa*. Unresolved siphonophore identifications were retained.

Church and colleagues contributed 151 libraries; Ahuja and colleagues (2024), 33; Ahuja and colleagues (2026), 21; and Mańko and colleagues, 8. All 151 *Physalia* libraries came from Church and colleagues. Their QC report and sample-status table identify 12 upstream exclusions: YPM-IZ-110432, YPM-IZ-110973 and

YPM-IZ-110632 for heavy Vibrionales signals; YPM-IZ-110574 for arthropod sequences; YPM-IZ-110826 and YPM-IZ-110827 for cross-sample contamination; and YPM-IZ-110474-2, YPM-IZ-110694-dry, YPM-IZ-106944, YPM-IZ-110879, YPM-IZ-111015 and YPM-IZ-111019 as replicates or duplicates. We retained all ten libraries they scored as moderate quality. We use *Physalia mikazuki* for the five specimens recorded as *Physalia* sp1 (cluster B2) in the original dataset, following its taxonomic description [5, 6]. For *Nanomia*, we matched specimen vouchers in the mitochondrial trees of Ahuja and colleagues (Fig 4 and S8 Fig) to library identifiers in their S8 Table [3]. This distinguishes *N. bijuga*, *N. cara*, *N. septata* and *Nanomia* sp. 1 in the sample table.

We reconciled collection metadata against specimen spreadsheets, museum catalogs, public BioSample records, publication supplements and author clarifications. Stable analysis identifiers link to separate voucher, BioSample, collecting-event and tissue-sample identifiers in the sample table. Sources, conflicting records and missing values are listed in the provenance and metadata-gap tables. All 213 specimens have coordinates, collection dates and a depth value; tissue remains unknown for 2 specimens. When only a depth interval was available, we assigned its midpoint while retaining both endpoints (14 specimens). One specimen uses the mean of its reported event depths. The 150 *Physalia* without a recorded depth were assigned 0 m to represent their surface habitat. Collection depth was not included in the regional models.

We subsampled inputs above 200 million pairs with BBTools (seed 20260829), then detected paired-end adapters and filtered reads with fastp 0.23.2 using default quality settings and disabling duplication evaluation [7]. Candidate screens used Kraken2 2.1.3 against the Standard database released on 4 September 2024 (`k2_standard_20240904`), Bracken 3.0.1 with a nominal read length of 150 and a ten-read threshold, sylph 0.9.0 with unknown-fraction estimation, and phyloFlash 3.4.2 against SILVA 138.1 [8–12]. The sylph databases were GTDB r220 c200 dbv1 and OceanDNA c200 v0.3 [13, 14]. We retrieved OceanDNA references from its official species-representatives archive [14]. The Kraken database is available from the versioned release archive, with a checksum manifest. Software environments and analysis settings are recorded in the environment files and workflow configurations.

To remove host reads, we used *Physalia physalis* assembly GCA_041430235.2 for all 151 *Physalia* libraries and *Nanomia septata* assembly GCA_048301705.1 for all ten *N. septata* libraries. Both *Halistemma rubrum* libraries used assembly GCA_987347065.1. The shared *Physalia* reference followed Church and colleagues, who reported 85–99% mapping across host lineages [1]. For *Nanomia*, a preliminary check mapped three million read pairs per tested library to *N. septata*; only *N. septata* libraries met the 80% properly paired alignment criterion for host-read removal. We retained siphonophore reads in the other *Nanomia* libraries. BWA-MEM alignment followed by SAMtools selection of flag 12 retained pairs with both mates unmapped [15, 16]. Assembly used all retained host-depleted pairs, or a seeded sample of up to 25 million trimmed pairs for libraries without host depletion (seed 20260901). We attempted assembly of all 213 libraries. Subsequent mapping used all retained host-depleted pairs or full trimmed reads up to the 200-million-pair cap.

#### S1.2 Reconstructing bacterial genomes

We reconstructed bacterial genomes from each specimen. We assembled each library separately with MEGAHIT 1.2.9, retaining contigs of at least 1,000 bp [17]. We grouped contigs into metagenome-assembled genomes (MAGs) with MetaBAT2 2.15, using coverage from reads mapped to the assembly, a 1,500-bp minimum contig length and seed 42 [18]. We estimated completeness and contamination with CheckM2 1.0.1. We reassessed MAGSP0031 with the same version after removing its palindromic duplication for deposit preparation. We retained genomes with at least 50% completeness and less than 10% contamination; those with at least 90% completeness and less than 5% contamination were termed near-complete. Bins lacking assessable DIAMOND annotations were ineligible [19]. We assigned domains with GTDB-Tk 2.4.1 and release r220 [13, 20]. Bins without prokaryotic marker placement were excluded. RNA-gene requirements for MIMAG high-quality status were not assessed [21].

We grouped similar genomes using skani 0.3.2, selecting the best-quality representative first. The quality score was completeness minus five times contamination. Clustering required at least 95% average nucleotide identity (ANI) and alignment covering at least 50% of either genome in a qualifying pair [22]. The retained 104 MAGs formed 32 clusters. Reference genomes identified by sylph at adjusted ANI of at least 95% were combined with MAG species representatives. The same grouping rule and quality score selected one representative per species.

We recorded agreement among genus names from different databases as additional support. We checked GTDB accessions against NCBI and obtained verified replacements for superseded accessions. References passed the MAG quality thresholds; OceanDNA genomes also required prokaryotic placement. Decoys were sampled with seed 20260927, excluding genomes within 95% ANI of candidates.

We reviewed every candidate bin using predicted proteins and translated 1,000-nt windows searched against Swiss-Prot 2026_03 with DIAMOND 2.2.0 (`-very-sensitive`, $E \leq 10^{-5}$). Protein-domain assignments required at least 50 aligned amino acids and agreement among resolved domains within 5% of the best bit score; ambiguous and unresolved matches remained separate. Discordant evidence was examined against the retained November 2023 nr database. Broader gene-domain evidence, marker support, coding density and assembly coverage were considered together. Known-origin proteins from all three domains served as assignment controls.

Contigs were also compared with the three host references and the excluded *Perkinsela*-related bins using minimap2 `-x asm20 -secondary=no -c`. We removed broad matches covering at least half a contig at 75%

identity when bacterial protein support was absent or limited to one Swiss-Prot gene/window and nr supplied no bacterial assignment. We also removed fragments with eukaryotic protein and translated-window support, no bacterial support, and either two eukaryotic nr assignments or one spanning at least 150 amino acids. Short isolated cross-domain matches in bacterial gene contexts did not force exclusion. The contig decisions and bin decisions record the evidence. Edited bins were reassessed with CheckM2 before clustering. Four bins with apparent prokaryotic marker placement instead showed *Perkinsela*-related eukaryotic identity and were excluded from the prokaryotic mapping catalog.

#### S1.3 Testing bacterial presence

To assess presence across specimens, we mapped reads with BWA-MEM using `-M -T60`; alignments were restricted to primary, properly paired records at MAPQ $\geq 30$, followed by duplicate removal. We measured genome coverage and read counts with CoverM, requiring identity $\geq 95\%$ and aligned read fraction $\geq 75\%$ [23]. Genome breadth was the fraction of catalog bases covered by retained reads. We checked whether MAGs were recovered by mapping reads from the libraries used to assemble them, and assessed 20 decoy genomes across all 213 libraries. The source-library comparison checks recovery through the mapping and filtering steps. Decoys, selected outside candidate genome clusters, check for gross spurious mapping. These checks do not calibrate discrimination among close relatives or sensitivity at low abundance. Real contaminants can pass the presence criteria, and genuine low-abundance associates can fall below them.

To flag reads concentrated in small parts of a genome, we compared observed breadth with that expected from uniform coverage. The ratio was $b/[1 - \exp(-150r/L)]$, where $b$ is breadth, $r$ is read count and $L$ is reference length. This calculation assumes 150-bp reads. We flagged ratios below 0.5 and retained these detections in the results. We aligned assemblies against catalog v3 using minimap2 `-x asm5 -c -secondary=no`. Assembly support required either a MAG from the same genome cluster in that specimen or at least 10 kb of catalog sequence aligned at 95% identity. We used the union of aligned catalog intervals, counting overlapping alignments once. The smaller assembly inputs used without host depletion can make assembly support harder to obtain.

#### S1.4 Metamycoplasmataceae relationships and 16S comparisons

We inferred a phylogeny using GTDB-Tk's bac120 alignment of 120 conserved bacterial marker proteins. The analysis included the seven recovered Metamycoplasmataceae genome representatives, 29 qualifying external host-associated genomes, GTDB Mycoplasmatales and Acholeplasmataceae. We analysed the 483-genome, 5,035-amino-acid alignment with IQ-TREE 3.0.1 [24]. ModelFinder selected LG+F+I+R10 by the Bayesian information criterion [25]. We conducted five independent maximum-likelihood searches and retained the tree with the highest likelihood. Each search included 1,000 SH-aLRT replicates [26, 27] and 1,000 ultrafast bootstrap replicates, with additional nearest-neighbour interchange optimization of bootstrap trees (`-bnni`) [28]. The reported support values come from the selected search. We rooted the displayed tree on Acholeplasmataceae. Support describes the concatenated-marker analysis; it does not measure agreement among individual markers. MAG 16S ribosomal RNA genes of at least 1,200 bases were available from MAGs in five of the seven clusters. We excluded a misbinned clade-B 16S copy from the MAGSP0005 bin WAM-Z97916 bin.2 before matching (contig decisions). Amplicon sequence variants (ASVs) from six Cnidarian Microbiome Database ASV libraries were matched across their full query length at 97% identity, retaining best matches and ties; studies were counted by publication [29].

Independent *Cassiopea* datasets were PRJNA1020388 (96 Illumina V3–V4 runs) and PRJNA1020446 (three PacBio HiFi runs) [30]. With VSEARCH 2.29.1 [31], we merged Illumina pairs allowing ten differences, removed the 341F/805R primer-length ends and retained 380–480-base sequences with at most one expected error and no ambiguous bases. Dereplication, UNOISE denoising with minimum abundance eight, and de novo chimera removal preceded matching to MAG 16S genes from 90% identity upward. HiFi reads were restricted to 1,300–1,700 bases.

#### S1.5 Identifying viruses

We searched assemblies for viruses with geNomad 1.12.0 and assessed their quality with CheckV against database v1.5 [32, 33]. Retained candidates required a viral hallmark gene and either length of at least 5 kb or estimated completeness of at least 50%. Direct terminal repeats (DTRs) alone did not establish viral identity. Sequences aligning across at least half their length to an available host reference were designated endogenous candidates (minimap2 `-x asm20 -secondary=no -c`; union of aligned query intervals, without an additional identity cutoff). The *N. septata* reference also supplied a congener comparison for reference-free *Nanomia*. Preplasmiviricota, Duplornaviricota, Negarnaviricota and Kitrinoviricota were also treated as endogenous candidates. The latter three are RNA-virus groups; their sequences in these DNA libraries could derive from integrated copies. This classification does not establish integration. The classification table records the assignments. We grouped sequences with CheckV into viral operational taxonomic units (vOTUs) at 95% ANI over 85% of the shorter sequence [34]. CheckV cellular-boundary calls were used to extract viral regions; separate regions on the same contig were retained separately. A second CheckV assessment confirmed that the retained regions lacked called cellular flanks and still met the hallmark and length/completeness criteria. The assembly-quality stage used CheckV 1.0.3; the vOTU stage used 1.1.1.

Associate and endogenous-candidate vOTUs were mapped competitively together, using the bacterial alignment filters above (primary proper pairs, MAPQ at least 30, duplicate removal, at least 95% read identity and at least 75% of each read aligned). The viral presence threshold was 75% breadth; shared near-identical viral modules can reduce uniquely mapped breadth and cause source-library non-detection. Viral negatives were not available.

#### S1.6 Identifying eukaryotes and interpreting their roles

The reference for the small-subunit ribosomal RNA gene (SSU; 18S in animals) combined SILVA NR99 with host SSUs assembled from the study specimens. Reads competed for placement against this combined reference. We aligned reads with minimap2 using `-ax sr -N 200 -p 0.99` and retained secondary alignments [35]. Both mates had to align for at least 100 bases. The pair's identity band was determined by the lower of its mates' best identities. Its lineage was the lowest common ancestor (LCA), the narrowest taxonomic group shared by both mates' qualifying matches. Cnidarian assignments and unresolved Metazoa or broader ancestral groups were excluded from read evidence. Insect, land-plant and non-human-mammal assignments provided background controls. Across the 205 libraries used for calibration, their maximum background was 36 pairs at 97% identity, so read-based acceptance required at least 37 pairs. A non-host assembled SSU also sufficed for acceptance, including when read counts fell below this threshold. We retained the read threshold for all 213 libraries and excluded specimens whose control counts exceeded the calibration range from biological analyses. Matches supported by two to 36 read pairs at 97% identity were retained as trace evidence. These were computational background controls, not extraction blanks.

Assembled evidence included phyloFlash SSUs and barrnap 0.9 18S segments of at least 500 bases on same-library contigs, classified against the same reference. Thus assembled support need not represent a complete eukaryotic genome or complete SSU. We checked assembled non-host sequences against NCBI, retaining broader identifications when a closest match could not resolve species. SILVA species labels were not reported. Identity $\geq 97\%$ allowed identification no finer than genus; 90–97% permitted family/order, and lower identities permitted class/phylum. An order-level LCA remains order-level even when identity permits a genus ceiling.

When accepted evidence identified both a broad group and a subgroup in the same specimen, the broader assignment was retained in the evidence table but not counted again. A trace-level subgroup did not replace an accepted broader assignment, and reads assigned broadly were not reassigned to subgroups. Ecological roles were assigned from taxonomic identity and known biology. Verified *Perkinsela*-like kinetoplastid matches were labelled endosymbionts of *Paramoeba*; unverified kinetoplastid evidence remained unassigned.

#### S1.7 Fish mitochondrial 12S screening

We searched all retained pairs from 163 host-depleted libraries and capped, quality-filtered full reads from the other 50 libraries. The September 2026 MitoFish snapshot supplied annotated 12S sequences [36]. For candidate recruitment, low-complexity bases were masked to N with DUST (window 64, level 20, linker 1). The same reference was searched independently with a conserved-sequence regex in both orientations, BBDuk exact 25-mers, Bowtie2 `-very-sensitive-local` and SortMeRNA [37, 38]. The execution guide and reference inventory retain commands, input selection and checksums.

Candidate pairs were compared by BLASTN with unmasked MitoFish 12S and the 3 September 2026 mitochondrial RefSeq release [39]. Strong evidence required an alignment of at least 100 nt at 95% identity, fish-only near-best matches within 5% of the best bit score, an adequately resolved hit-list tail, and no strong competing non-fish match from the other mate. RefSeq identity was established from accession-linked GenBank annotations. Whole-mitochondrial alignments can extend beyond 12S; the pair table records overlap with the marker, both mates' matches and reference ambiguity. Synthetic fish and non-fish controls checked recruitment and competitive classification. Exact paired-sequence hashes assessed duplication. The 37-pair SSU presence criterion was not applied to this different marker.

#### S1.8 Direct comparison with published trematode sequences

We retrieved all 108 nucleotide sequences deposited by Waki and colleagues and linked to their publication record (26 28S, 11 ITS2-containing amplicons and 71 COI sequences), together with 113 external reference sequences cited in their supplementary distance matrices [40]. Reference metadata retain accession versions, host and life-stage annotations; retrieval records identify source files and checksums. The ITS2-containing records span 5.8S, ITS2 and 28S but do not annotate internal boundaries; reported identities therefore apply to the aligned amplicon rather than to ITS2 alone.

We searched all 221 references against assembled contigs from the eight specimens with trematode SSUs using BLAST+ 2.5.0 [39]. The nucleotide search used task `blastn`, word size 11, DUST masking, an expectation threshold of $10^{-10}$ and retention of up to one million target sequences; all qualifying local alignments were retained. Alignments of at least 200 bases were tabulated as homologous-region candidates, without a species-assignment threshold. Competing references were compared over shared contig positions when each covered at least 80% of the focal alignment interval. This comparison excludes insertions relative to the contig and retains reference ambiguity separately. The two COI regions represented in the reference collection were compared separately when they did not overlap. The accepted parasite SSUs were also aligned to these contigs to distinguish physical linkage from recovery in the same specimen.

#### S1.9 Testing geographic and sequencing-batch associations

Flowcell identifiers were parsed from FASTQ headers, excluding lane fields from `instrument:run:flowcell:lane`; multiple lanes of one flowcell remain eligible. Inference used 140 retained single-flowcell libraries, including 122 *Physalia*. Multi-flowcell libraries remain in descriptive results.

For genomes with at least three eligible presences, flowcell concentration was $\sum_f n_f^2/N$, calculated using presences in host-taxon-by-region strata with at least two physical flowcells. Flowcell labels were permuted 10,000 times within strata, with seed 20260929. Insufficient eligible counts, nonpermutable presences and constant statistics were explicitly untestable. One-sided empirical probabilities used $(e+1)/(B+1)$, where $e$ permutations met or exceeded the observed statistic and $B$ is the number of permutations. Benjamini–Hochberg adjustment used only testable genomes. The cohort and *Nanomia*-only tests formed separate families.

Regional models were restricted to Physalia and required at least two regions on two flowcells each and a flowcell spanning regions. The planned mixed models fitted unreliably; their convergence diagnostics are retained with the statistical results. We instead used the logistic-deviance gain beyond $\log_{10}$ mapping-input pairs, with 9,999 region-label permutations within flowcell and seed 20260930. Jaccard PERMANOVA entered log-transformed input read-pair count before region and restricted permutations within flowcell. Empty communities were excluded, with denominators recorded; identifiability was reassessed on that subset. We ran these analyses in R 4.4.3 with lme4 2.0-6 and vegan 2.7-5.

The complete incidence and statistical analyses were repeated at 5% and 20% bacterial breadth, including threshold-specific selection of taxa with ten eligible *Physalia* presences. A separate fixed-catalog sensitivity experiment tested host handling in five *Physalia* and five *N. septata* libraries, selected across retained-input-depth quintiles (seed 20260930). Capping was tested in eight libraries above the cap, split equally between libraries processed with and without host depletion (seed 20261001). Both sides of the cap comparison used full trimmed reads to isolate capping. These comparisons used the earlier catalog v2 and the specified library subsets. Their scope differs from the full-cohort results, which use catalog v3. We also counted candidate genomes identified only in one study to describe how reference selection depended on that study. This comparison held the catalog fixed and did not test read reassignment after removing references.

#### S1.10 Tissue, depth and targeted host–region comparisons

We linked accepted detections to the curated sample table by library identifier, retaining specimens without detections. Summaries counted positive specimens separately from genome–specimen, viral-group–specimen and broad eukaryotic-group detections. Eukaryotic summaries used the counting criteria to avoid counting ancestor and descendant assignments twice. Mapped presence and same-specimen assembly support were summarized separately. A zero denotes a result below the detection threshold.

Depth displays retained recorded points and interval endpoints. *Physalia* surface values were summarized separately from other hosts. Depth groups distinguished intervals wholly within 0–20 m, intervals crossing 20 m, and collections deeper than 20 m; the grouping used interval bounds even where a midpoint was assigned. Tissue summaries retained the source descriptions, with unrecorded tissue explicitly unknown. We compared detections of a Metamycoplasmataceae bacterial lineage labelled DT-68 in the Genome Taxonomy Database (GTDB) across retained *Nanomia* and *Resomia* specimens. Regional incidence for three previously identified Metamycoplasmataceae genome clusters and *Vibrio* was tabulated for all *Physalia* and recorded tentacle samples, with an additional within-*P. utriculus* comparison. Single-flowcell subsets were tabulated separately (source tables).

We compared associate detections across tissues and collection depths using all retained specimens. We assessed the replication and overlap of tissue, host identity, locality and flowcell before attempting statistical tests. Life-stage labels were not analysed because they were inconsistently recorded. The depth-design table and contrast-design table record these checks.

For targeted community comparisons, we selected tentacles of *P. utriculus* from the central, northwestern and southwestern Pacific, and tentacles of *P. megalista*, *P. minuta* and *P. utriculus* from the southwestern Pacific. Both comparisons used the single-flowcell subset and excluded empty bacterial profiles from Jaccard distances. Marginal PERMANOVA tested the group effect after log-transformed actual mapping input pairs and flowcell, with 9,999 permutations restricted within flowcell. We required a full-rank group contrast and at least 20 distinct within-flowcell group-label allocations. An additional *P. utriculus* contrast including the western Indian Ocean failed the rank check; Eastern Indian Ocean libraries all spanned multiple flowcells. A matched tissue comparison within central Pacific *P. utriculus* lacked sufficient distinct permutations.

These post hoc comparisons used the primary, 5% and 20% breadth definitions. Benjamini–Hochberg adjustment covered estimable contrasts within each threshold. Exact subsets, settings and software versions are provided with the analysis tables and provenance. Fish read-pair counts in tissue summaries came from the separate 12S screen; they were not pooled with genome-target counts.

#### S1.11 Human-read assessment and bacterial-background comparison

To assess human DNA and its association with bacterial background, we selected up to one million pairs per library from equally allocated prefixes of its raw source files. These convenience subsets may differ from the complete libraries. After fastp filtering, Bowtie2 end-to-end alignment against the locally indexed UCSC hg38 genome used `-very-sensitive -no-mixed -no-discordant`, a fixed maximum fragment length of 1,000 bp and its standard score range [37]. Both mates had to be primary, properly paired, unclipped alignments of at least 100 nt, with at least 99% identity and MAPQ 30; low-complexity sequences were excluded. Exact sequence-pair duplication and occupied 1-Mb windows were summarized separately. The human-screen methods provide the reference identities and complete filtering settings.

Known-origin controls used 10,000 simulated pairs from each of hg38 and the three siphonophore references, with 150-nt reads, 350-bp fragments and one substitution per mate. These measure recovery and cross-mapping under those conditions; they do not establish specificity against all nonhuman genomes or other read-length distributions.

We compared up to 500 distinct candidate pairs per library with the local 2023 NCBI nt snapshot using megablast [39]. Matches required at least 98% identity across 95% of the query; near-best matches had bit scores within 5% of the best. Each mate was assessed separately, and a pair was called human-only among near-best matches only when both mates met that condition. Searches reaching the 100-subject cap, unknown taxonomy and incomplete mate confirmation remained unresolved. We kept counts of read pairs matching the human reference genome separate from these taxonomic classifications.

We compared human-compatible read pairs per million screened pairs with *C. acnes* BAC00019 breadth in all libraries (Fig H). The specimen comparison retains the underlying counts. We set the exclusion thresholds after inspecting gaps in these distributions: *C. acnes* breadth at least 50%, human-compatible pairs at least 0.05%, or SSU controls above the calibration range. The excluded-library table records the criteria met by NA29, NA31, D666-D10 and T676-SS3.

In retained libraries, we set aside skin-associated *Cutibacterium* and *Lawsonella* [41, 42]; the reagent-associated groups *Bradyrhizobium*, *Mesorhizobium*, *Methylobacterium*, *Pelomonas*, *Pseudomonas*, *Afipia*, *Sphingomonas*, *Acidovorax* and *Acinetobacter* [43]; the *Serratia* clone MAGSP0001; and the *C. elegans* SSUs in TMAG-K5632 and TMAG-K5633, both scored as moderate quality by Church and colleagues [1]. The contaminant table records these assignments. Detections remain flagged in the full evidence tables but do not contribute to biological counts, incidence or community tests. No extraction blanks were available. All ten moderate-quality *Physalia* libraries were retained.

#### S1.12 Analysis code and sequence deposits

We ran the analyses with Snakemake [44]. Analysis and figure-generation code, configuration and provenance are available at https://github.com/caseywdunn/siph-associates, with the earlier v0.5.0 code release archived at https://doi.org/10.5281/zenodo.23441461. Submission of the reconstructed sequences to NCBI is pending. The repository's submission guide links preparation files and records the current status.

### S2 Supplementary results and sensitivity analyses

#### S2.1 Sampling and genome recovery

Each of the 213 siphonophore specimens was represented by one sequencing library. Of these specimens, 207 were assigned to 37 named siphonophore taxa; the remaining 6 had broader or provisional identifications. Sampling was greatest for the surface-dwelling *Physalia* (151 specimens) and *Nanomia* (22 specimens across two studies; Fig A). 31 host identifications were represented by one specimen each.

The samples span the sea surface, the water column and the seafloor. The epibenthic rhodaliids were represented by one *Stephalia dilata* collected at 1,349 m. Collection depths were recorded or assigned for all 213 specimens, including 151 surface-dwelling *Physalia* (Fig B shows the 209 retained specimens). Assembly was attempted for every library.

Protein-domain and contig review retained 104 bacterial MAGs in 32 clusters; 55 met the near-complete criteria. The catalog combines 24 MAG-derived and 454 reference-derived representatives, with 20 decoys. Catalog representatives differ from recovered MAG clusters because some clusters are represented by a public reference after joint dereplication. We recovered MAGs with and without host depletion: 96 came from host-depleted libraries and 8 from libraries assembled without host depletion.

No decoy genome met the presence criterion at any of the three tested breadth thresholds. The primary 507 bacterial genome–library detections included 294 with same-specimen assembly support and 32 with the flag for reads concentrated in small genome regions. A further 1,151 pairs were trace evidence. Among accepted pairs, 285 of 467 after host depletion and 9 of 40 without host depletion had assembly support. These proportions also reflect different assembly input strategies.

#### S2.2 Bacterial detections

Bacteria contributed 507 detections of 166 representatives in 131 libraries, including 294 with assembly support (microbial detection table).

Metamycoplasmataceae contributed 138 detections of 6 representatives in 76 libraries, including detections with assembly support. The GTDB genus *Tyloplasma* was represented by MAGSP0033 (BAC00459), in Hepatoplasmataceae. This genome was recovered from YPM-IZ-111016 and also detected in YPM-IZ-111017; it lies outside Metamycoplasmataceae. *Vibrio* reads matched 38 reference genomes across 25 libraries, giving 106 genome–library matches. The recovered *Vibrio* MAGs were MAGSP0014, assigned to *V. caribbeanicus* and meeting the near-complete criteria, and MAGSP0023, assigned to *V. alginolyticus*.

#### S2.3 Metamycoplasmataceae relationships and host distributions

Clades A and B were detected only in *Physalia* after sample exclusions; DT-68 occurred in seven *N. septata* and two *Resomia ornicephala*. MAGSP0031 formed a divergent sister lineage to clade B. MAGSP0012, assigned to *Mycoplasma_K*, was recovered only from NA31, a specimen excluded from biological analyses, and could have arrived with prey. It remains in the phylogenetic comparison.

MAGSP0031 was detected in three retained *P. utriculus* specimens. Its catalog version contains a palindromic duplication of about 80 kb. This region did not contribute uniquely mapped reads and therefore lowered the measured breadth; we did not repeat read mapping. Removing the duplication yielded 693,250 bases, with CheckM2 estimates of 87.49% completeness and 0.00% contamination (corrected-genome report).

The smallest clade containing all seven siphonophore-associated Metamycoplasmataceae genomes had 21 tips representing 20 distinct assemblies: one bacterial genome recovered from an octocoral occurred as both an external input and a GTDB reference. The surrounding tree included 483 tips, including 29 external host-associated genomes retained from 32 candidates. The recovered bacteria group with relatives from other marine animals, although the clade-B/DT-68 sister relationship was weakly supported (SH-aLRT 64.2%, ultrafast bootstrap 71%).

MAGs in five of the seven clusters contained suitable assembled 16S genes for external marker comparisons. Matching ASVs occurred in 98 samples from 23 publications in the Cnidarian Microbiome Database. ASVs identical to clade-B genes occurred in the coral *Astroides calycularis* and the sponge *Aplysina aerophoba*. These short-marker matches include related lineages and cannot establish membership in the same genome cluster. No *Cassiopea* denoised Illumina variant or length-filtered HiFi read matched the available siphonophore genes at 90% identity or higher. This non-detection is restricted to the queried genes and sampled datasets; the HiFi comparison included 38,067 length-filtered reads.

#### S2.4 Regional distributions and sensitivity to detection thresholds

Within *Physalia*, three Metamycoplasmataceae genome clusters varied significantly among sampling regions: MAGSP0005 in clade A, and MAGSP0007 and MAGSP0010 in clade B. A total of 5 bacterial genomes had enough detections for these tests. The tests accounted for the number of reads available for microbial detection and compared regions within physical sequencing flowcells.

All three were frequent in the southwestern Pacific and rare in the central Pacific. Among the 50 southwestern Pacific specimens, MAGSP0005 occurred in 24, MAGSP0007 in 14 and MAGSP0010 in 24. These specimens came from New Zealand (23), New South Wales (11), Tasmania (eight), the Tasman Sea (five) and Queensland (three). By contrast, each genome cluster occurred in only one of 23 central Pacific specimens, principally from Hawai'i.

The Indian Ocean was represented by 17 specimens. Among the 12 from Western Australia, MAGSP0005 occurred in 11 and MAGSP0010 in nine. Among the five from South Africa, MAGSP0005 occurred in four and MAGSP0010 in all five. These counts retain all libraries, including those sequenced on multiple flowcells. The statistical tests used the single-flowcell subset and assessed differences among all regions together.

The *Physalia* per-taxon comparison included 11 geographic categories: central, northwestern, southeastern and southwestern Pacific; northeastern, northwestern and southwestern Atlantic; eastern and western Indian Ocean; and the gulfs of California and Mexico. The 122 eligible libraries represented all 11 categories. At the primary threshold, all seven eligible Gulf of Mexico libraries lacked accepted bacterial detections. The community comparison comprised 88 nonempty libraries. The comparison distinguished named regions within and among ocean basins.

Among 40 genomes with at least three total accepted presences, 26 had testable concentration statistics; the other genomes lacked sufficient eligible or permutable presences, or had a constant statistic. None of the 26 tests passed the 5% false-discovery threshold. Neither of the two genomes with sufficient total presences in the pooled *Nanomia* analysis had enough eligible, permutable detections for a test. These results do not establish freedom from contamination, particularly without extraction blanks.

Descriptive bacterial detections decreased from 750 pairs (251 targets, 143 libraries) at 5% breadth to 507 (166 targets, 131 libraries) at 10%, and 342 (112 targets, 121 libraries) at 20% (Table A). The eligible *Physalia* denominator remained 122. At 5%, 97 nonempty libraries entered PERMANOVA; ocean region explained 22.2% of variation ($P \leq 0.0002$), and 7 of 9 eligible genomes had regional permutation-test $q < 0.05$. At 10%, exclusion of empty communities left 88 libraries for PERMANOVA; ocean region explained 19.8% of Jaccard variation after input read count ($P \leq 0.0002$). 3 of 5 eligible genomes had regional permutation-test $q < 0.05$. At 20%, the community analysis included 81 libraries, and region explained 22.6% ($P \leq 0.0002$); 2 of 4 tested genomes passed the adjusted significance threshold. The fitted mixed models retained convergence warnings; their likelihood-ratio probabilities were not substituted for the permutation results. Community-level regional association persisted across all three thresholds, while the tested taxon sets and some individual conclusions changed.

#### S2.5 Targeted comparisons of hosts, regions and tissues

The southwestern Pacific host comparison included 30 nonempty bacterial profiles from *P. megalista*, *P. minuta* and *P. utriculus* tentacles. Host identity accounted for 13.3% of variation with read effort and flowcell as covariates and permutations within flowcells ($P \leq 0.0002$; $q = 0.0002$). The effect persisted at 5% breadth ($R^2 = 12.6\%$, $P \leq 0.0002$) and 20% breadth ($R^2 = 9.6\%$, $P = 0.0233$, $q = 0.0466$).

Within *P. utriculus*, the three Pacific regions contributed 31 nonempty tentacle profiles at the primary threshold ($R^2 = 6.1\%$, $P = 0.2916$). A regional difference was detected only at 5% breadth ($P = 0.0068$), with no significant difference at 20% ($P = 0.1269$). The community tests include all thresholds and exclusions. Incidence summaries retain the empty profiles and libraries ineligible for inference (Fig F).

Tissue categories and collection depths were strongly associated with host and locality (Fig G). Alternative tissues within comparable hosts and regions were too sparsely replicated for an independent tissue test. The two positive zooid samples, YPM-IZ-110906 and YPM-IZ-110907, contributed 19 of the 32 qualifying fish 12S pairs, compared with 13 across the positive tentacles. Differences in read effort and the small number of zooid specimens preclude treating these counts as a measure of tissue-specific prey abundance. The two positive zooid samples came from the same northwestern Hawaiian locality and shared sequencing lanes.

#### S2.6 Sampled tissues and within-tissue regional comparisons

*Physalia* samples included 142 tentacles, with smaller numbers of zooid, whole-body and corm samples and 2 of unknown tissue (Fig D). The retained samples from other hosts were siphosomes.

The southwestern-versus-central Pacific contrast for MAGSP0005, MAGSP0007 and MAGSP0010 remained visible within recorded tentacle samples, including the subset of *P. utriculus* tentacles. The regional tables give positive counts and denominators for all *Physalia*, tentacle-only and single-flowcell subsets. The host-specific comparison retains differences in collection history and sequencing batch.

#### S2.7 A Metamycoplasmataceae lineage occurs in specimens collected at greater depths

Collection depths spanned the surface habitat of *Physalia*, the water column and the seafloor (Fig B). Host identities and sampling histories differed across this range.

The comparison of the bacterial lineage DT-68 included the retained *Nanomia* and both *Resomia ornicephala* specimens (Fig C). All seven California *N. septata*, collected at 217–662.29 m, carried DT-68, whereas none of the three Washington specimens, collected at 0–20 m, did. California detections included both read-only and assembly-supported evidence. Both *Resomia* specimens, collected at 235 and 286.25 m, had assembly-supported detections. DT-68 was not detected in retained *N. bijuga* or *N. cara* specimens, all collected at 0–20 m. These compare different host identifications. All ten *N. septata* were siphosome samples from the same source study. Locality and depth remain confounded.

#### S2.8 Viral detections

The associate-virus catalog contained 401 viral operational taxonomic units (vOTUs). Of these, 256 met the read-based presence threshold in at least one of 119 specimens, giving 478 detections. For 297, a sequence from that viral group was also assembled from the same specimen. Most groups belonged to Uroviricota, which contains tailed bacteriophages; other groups included Cressdnaviricota and larger DNA viruses. A separate catalog contained 163 endogenous candidates, including sequences assigned to the RNA-virus groups Negarnaviricota and Kitrinoviricota that could represent integrated copies.

#### S2.9 Eukaryotic diversity

Using SSU markers, we detected 15 broad groups of non-siphonophore eukaryotes in 38 specimens. Counting each group once per specimen gave 57 detections, of which 47 were assembly-supported; 40 had both reads and assembly (main-text Fig 4). Animal detections included parasitic flatworms in 11 specimens, chordates in four, arthropods in three, molluscs in two and a chaetognath in one. Other recovered groups included amoebae, kinetoplastids, fungi, dinoflagellates, algae and ciliates.

The eukaryote evidence table contains 58 accepted assignments, including overlapping classifications counted only at the more specific level.

Four related genomic bins showed strong *Perkinsela* affinity. These bins initially received a prokaryotic placement from GTDB-Tk, but protein-level review assigned 2,518 of 3,615 predicted proteins in their representative MAGSP0026 to eukaryotes. All 29 hits to the prokaryotic marker panel matched *Perkinsela* proteins. We excluded these bins from prokaryotic counts and retained them for further eukaryotic characterization (domain-review tables).

#### S2.10 Fish markers and reference ambiguity

The fish 12S screen examined 9,313,145,671 pairs. BBDuk recovered all 32 strong matches; Bowtie2 and SortMeRNA each recovered 31, and the motif regex recovered 7. All 32 paired sequences were distinct and occurred in 8 *Physalia* libraries. Exact sequence uniqueness does not establish independent original DNA molecules or a number of prey individuals. Near-tied reference names remain identification alternatives, and a negative library does not establish that its host never feeds on fish.

Reference affinities included anchovies (*Engraulis*), *Beryx* and carangids, with near-tied matches limiting species identification. No additional qualifying matches occurred in host-depleted *N. septata* or *H. rubrum* reads, or in the 50 other libraries, which contributed 8,893,919,800 quality-filtered pairs. Near-tied reference matches were retained as identification alternatives (pair-level evidence).

Fish-assigned reads occurred in four *Physalia* libraries: *P. utriculus* WAM-Z97917 from New South Wales and YPM-IZ-110907 from the northwestern Hawaiian Islands, and *P. megalista* YPM-IZ-111156 and YPM-IZ-111158 from the Tasman Sea. Each contained reads assigned to Teleostei. Only YPM-IZ-110907 yielded an assembled fish SSU, 1,834 bases long. Its two highest-scoring archived NCBI matches were predicted *Centroberyx gerrardi* 18S sequences XR_013504110 and XR_013507526, both 99.95% identical over all 1,834 bases. The third retained hit, XR_013507952, was also labelled *C. gerrardi* and had 99.89% identity. These searches used `core_nt` on 30 September 2026. The retained hits represent multiple 18S copies from one named reference taxon. We report this close reference affinity and retain Teleostei as the accepted assignment for all four specimens.

The presence threshold was applied to the broad Chordata group, combining Teleostei reads with reads assigned to broader chordate lineages. The evidence table therefore reports the combined chordate read count. The three read-only specimens had Teleostei as their most frequent exact read-level assignment, but no assembled fish sequence for independent reference comparison.

#### S2.11 Flatworm markers and comparisons with published parasites

The 11 parasitic flatworm detections contained 12 assembled SSUs: trematodes in eight specimens and cestodes in four, with both in *P. physalis* YPM-IZ-110436 from Florida. The broad evidence table counts Platyhelminthes once per specimen and uses one representative lineage label; its eight trematode-labelled and three cestode-labelled rows therefore conceal this overlap. Counting each broad group once per specimen gives 11 parasitic flatworm detections despite the two parasite groups in one specimen.

Three 2,009-base trematode SSUs from *P. physalis* YPM-IZ-110436, *P. utriculus* YPM-IZ-110882 and *P. mikazuki* YPM-IZ-110931 matched *Dinurus longisinus* AJ287501 at 99.90–100% identity over 1,967 aligned bases. A 1,338-base SSU from *P. minuta* YPM-IZ-110557 matched *Prodistomum orientale* LC850120 at 99.16% over 1,313 bases, with *Lepocreadium trulla* and *Preptetos caballeroi* nearly as similar (98.94% over 1,315 bases). The *Preptetos* affinities in *P. physalis* YPM-IZ-110631, YPM-IZ-110694 and YPM-IZ-110846 likewise record the nearest archived references.

The 2,135-base cestode SSU from *P. minuta* YPM-IZ-111243 matched *Echinophallus wageneri* DQ925311 at 99.95% over 2,129 bases. Cestode SSUs from *P. physalis* YPM-IZ-110436 and *P. utriculus* YPM-IZ-110777 had competing *Crossobothrium*, *Marsupiobothrium* and *Scyphophyllidium* matches, with different alignment lengths and identities of 96.38–97.68%. The fourth cestode-positive specimen, *P. utriculus* YPM-IZ-110460, had nearly tied *Scyphophyllidium* and *Marsupiobothrium* scores. These SSU comparisons do not separate those three reference groups.

Damian-Serrano and colleagues used six short 18S amplicons; their classifications corroborate trematodes in *Physalia* and cestodes in other siphonophores [45]. We did not directly align their amplicons with the assembled SSUs. Waki and colleagues used 28S, ITS2 and COI to identify trematodes including *D. tornatus* and *P. orientale* in *P. utriculus* [40]. Our closest *P. orientale* 18S reference, LC850120, is an independent record rather than a sequence from their *Physalia* specimens. The additional markers below provide the direct comparison with Waki's records.

In *P. utriculus* YPM-IZ-110882 from Hawai'i, a contig matched the *D. tornatus* ITS2-containing reference LC889198.1 at 100% over 651 bases. In *P. mikazuki* YPM-IZ-110931 from Japan, the corresponding match was 99.55% over 665 bases. Separate COI contigs matched *D. tornatus* LC889235.1 and LC889234.1 at 98.15% over 867 bases and 98.61% over 863 bases, respectively. Both COI references were from metacercariae collected from *Physalia*. The ITS2-containing reference LC889198.1 was from a fish adult; the two contigs also matched the *Physalia*-derived larval reference LC889201.1 at 99.84% over 645 bases and 99.39% over 656 bases. These nuclear and mitochondrial matches support a *D. tornatus*-like interpretation. In Florida *P. physalis* YPM-IZ-110436, the ITS2-containing region instead matched *D. barbatus* LC889203.1 at 99.85% over 673 bases, compared with 99.10% over 665 bases for *D. tornatus* LC889198.1. The *D. barbatus* reference was from an adult in *Coryphaena hippurus*. No comparably close COI match was recovered for this specimen. These identities describe individual local alignments; the source table additionally compares competitors over shared contig positions.

In northeastern US *P. physalis* YPM-IZ-110631 and YPM-IZ-110694, 28S matches to *Prodistomum* Type 4 LC889183.1 were 99.39% over 1,149 bases and 99.14% over 1,167 bases, respectively. This reference was from a metacercaria in *Physalia*. The corresponding *P. orientale* reference matches were 97.2–97.5% identical; other *Prodistomum* references also matched closely. COI sequences in these two specimens were only about 87–88% identical to Type 4. These markers place the sequences among *Prodistomum* relatives, with COI divergence from Waki's Type 4 reference. The remaining three trematode-positive specimens (YPM-IZ-104464, YPM-IZ-110557 and YPM-IZ-110846) had no comparably close matches in this targeted reference set. Short or divergent matches remain in the full search results and were not converted to named parasite detections.

None of the additional marker contigs could be linked physically to an accepted SSU by an alignment covering at least 80% of the SSU at 97% identity. Multiple parasites may therefore contribute different markers within a specimen. *D. longisinus* was represented only by 28S in the comparison set, so the original closest 18S label does not distinguish it from the *Dinurus* alternatives above. The supporting `tables/waki_comparison/` files retain specimen, marker, accession, alignment length, identity, coverage, competing references and linkage status.

One Azygiida assignment matched accession Z23111, which retains a *Daphnia galeata* label despite its SILVA trematode placement. The corresponding archived NCBI search also closely matched *Accacoelium contortum*, supporting retention of the broad trematode assignment. This documented reference-label discrepancy was not interpreted as evidence for cladoceran prey or a more specific parasite identification.

#### S2.12 Support for initial read-screen assignments

Initial read screens identified candidates whose presence we then assessed by competitive mapping and assembly. We estimated genus and species abundances with Bracken from Kraken2 classifications, whereas phyloFlash SSU assignments included different taxonomic levels, siphonophores and unresolved groups. We compared these candidates with accepted detections by reference or group (screen-summary tables; Table B).

In the original screen of 213 libraries, sylph identified 505 matching reference genomes in 119 specimens. These included 355 GTDB and 150 OceanDNA genomes, with 977 genome–library matches in total. Genus corroboration was assessed for the GTDB subset, using Bracken support of at least ten estimated reads or phyloFlash support of at least one read. These permissive corroboration criteria were distinct from the final presence thresholds. The 69 GTDB genomes without same-genus corroboration included uncultured marine lineages with placeholder names or differing representation across reference databases. Lack of name agreement was therefore not used to reject sylph assignments.

For comparison with final mapping, references selected by sylph were represented by their retained catalog species representative and counted once per library. Excluded references were omitted, and references within the same genome cluster were counted together. Among retained specimens and taxa, competitive mapping supported 230 of 745 candidate genome–library pairs, including 97 with assembly support. The remaining candidates lacked sufficient coverage; low abundance and reference divergence can contribute to this outcome. Of the 515 candidate genome–library pairs below the primary presence threshold, 464 retained trace evidence and 51 had no graded support. The initial screens used quality-filtered reads before host depletion, whereas final mapping followed the specified host-read removal and competitive alignment filters. These differences, reference representation and the absence of an organismal ground truth preclude interpreting the rejection fraction as a measured false-positive rate for sylph, Kraken2 or Bracken.

The 507 accepted bacterial genome–library detections comprised 230 identified by sylph in the same library, 28 whose genome representative matched sylph references only in other libraries, and 249 involving the 24 representatives with no sylph match anywhere in the cohort. The last group included all 6 focal Metamycoplasmataceae representatives (138 detections in 76 specimens). Broad *Mycoplasma*-related labels were nevertheless present in Bracken and phyloFlash in all 76 Metamycoplasmataceae-positive libraries, using the permissive support thresholds above. The labels were identified by `mycoplasm` in Bracken genus names or phyloFlash SSU lineages, allowing for taxonomic differences among reference databases. Support was at the broad group level (accepted-detection overlap and Mycoplasma-related support tables).

The phyloFlash comparison used retained paired-read assignments at $\geq 97\%$ identity before and after host-aware competitive remapping. Both summaries required at least 100 aligned bases from each mate; the later assignment also considered near-tied reference matches and their lowest common ancestor. The 37-pair threshold was calibrated from a maximum of 36 control pairs in 205 libraries and applied throughout the cohort. The background-control table identifies specimens exceeding calibration; those specimens were excluded from biological analyses. Many initial insect assignments came from misclassified siphonophore reads. Initial insect SSU matches occurred in 186 libraries but remained in only 23 after competitive comparison with siphonophore SSUs. Conserved siphonophore reads had matched non-host references. After competitive comparison with host references, accepted detections still included SSUs from both phyloFlash and general assembly.

Among the 57 accepted nonredundant eukaryotic detections, 46 had a phyloFlash assembly and 12 also had an 18S sequence from the general assembly workflow. An additional kinetoplastid detection in YPM-IZ-111009 had an 18S sequence only from the general assembly. Its 17 supporting read pairs fell below the 37-pair threshold, so the assembled sequence supplied the confirming evidence. The sequence's closest verified NCBI match was a *Perkinsela*-like organism associated with *Paramoeba invadens.*

#### S2.13 Sensitivity to host-read removal, read count and reference selection

A comparison that held the bacterial and viral reference catalogs constant (catalog v2) while changing host-read removal comprised five *Physalia* and five *N. septata* libraries. It yielded 22 bacterial presences after host depletion and 24 using full trimmed reads. Twenty-one pairs agreed, one occurred only after host depletion, and three occurred only with full reads; four libraries changed. In the combined viral catalog, 22 presences after depletion compared with 53 using full reads: all 22 were retained and 31 additional vOTU–library pairs appeared. Every one of the ten libraries gained viral detections. These counts include both associate vOTUs and endogenous candidates.

Across eight libraries above the input cap, bacterial presences increased from 34 with capped full reads to 49 with uncapped full reads, with no losses. Twelve of the 15 additions occurred in Church2025 library YPM-IZ-111839, two in Ahuja2026 NA38 and one in Church2025 YPM-IZ-110436. In the combined viral catalog, uncapping added one detection in Ahuja2024 CWD3, increasing the total from 95 to 96, again without losses. Most of the additional bacterial detections came from a single library.

The nomination-dependence table records the reference genomes identified exclusively in each of the four studies and detections dependent on those references. MAG recovery supplied 24 representatives with no sylph match, accounting for 249 detections. These comparisons describe each study's contribution to the shared catalog; changes in competitive read allocation after removing a study were not tested.

##### S2.14 Human-compatible reads and contamination criteria

The controls retained 89.87% of quality-filtered simulated human pairs and no stringent matches among 29,355 quality-filtered siphonophore pairs. Of 483,068 read pairs initially mapped to the human reference genome, 4,563 passed stringent filters. These occurred in 178 of the 213 libraries screened. The largest count was 1,058 in NA31; T676-SS3 had 937.

Among 3,384 candidate pairs selected for nt comparison after within-library sequence deduplication, 2,060 had only human near-best matches for both mates and 1,241 also had alternative matches, including primate and synthetic-construct references. Confirmation was incomplete for 29 pairs; 53 were unresolved because of the hit cap or unknown taxonomy. No pair had conflicting mate assignments. The 1 pair lacking human near-best support matched a synthetic-construct reference. These classifications describe the selected candidate subset and available references (per-library summaries and controls).

The human-read and *C. acnes* distributions used to select the four excluded libraries are shown in Fig H. The largest *C. acnes* breadths occurred in NA29, NA31 and D666-D10; T676-SS3 exceeded both the human-read and SSU-control criteria. Contaminant detections in the remaining libraries were retained as flagged rows in the evidence tables and omitted from biological summaries.

#### S3 Supplementary figures and tables

| Breadth (%) | Detections | Genomes | Specimens | Community $n$ | Region $R^2$ | Region $P$ | Taxa with $q < 0.05$ |
|---|---|---|---|---|---|---|---|
| 5 | 750 | 251 | 143 | 97 | 0.222 | $\leq 0.0002$ | 7/9 |
| 10 | 507 | 166 | 131 | 88 | 0.198 | $\leq 0.0002$ | 3/5 |
| 20 | 342 | 112 | 121 | 81 | 0.226 | $\leq 0.0002$ | 2/4 |

**Table A.** Sensitivity of bacterial results to the fraction of each genome covered by reads (breadth). Detections count each genome once per specimen. Descriptive columns include the 209 retained libraries and exclude contaminant taxa and decoy genomes; every presence additionally requires at least 100 reads. The primary breadth threshold is 10%. Flowcell-based *Physalia* inference uses 122 single-flowcell libraries. Community $n$ excludes empty communities at each threshold; region statistics are sequential Jaccard PERMANOVA effects after log-transformed input read-pair count, with 9,999 within-flowcell permutations. The last column gives significant per-taxon regional permutation tests divided by the number tested; each taxon must occur in at least ten eligible libraries, and Benjamini–Hochberg adjustment is applied separately at each threshold. Changing community membership and tested taxa must be considered when comparing thresholds.

| Data directory | Contents and interpretation |
|---|---|
| Cohort and detections | Library metadata, bacterial and viral detections, and all eukaryotic evidence rows, including contamination flags, viral classification, ancestor assignments, counting flags and taxonomic resolution. |
| Overview | Study and domain summaries, microbial detection frequencies by host, display-group assignments and the eukaryote matrix with host and specimen labels. |
| Metadata audit | Collection-metadata additions, source conflicts, coverage, review decisions and retrieval provenance. Depth intervals retain both endpoints. |
| Collection context | Detections joined to collection metadata for the 209 retained specimens; depth, host and tissue summaries; DT-68 specimen context; and regional incidence within *Physalia* tissue and host subsets. |
| Genome and viral catalogs | MAG quality, species membership and mapping-catalog composition. The reviewed viral manifests record associate and endogenous-candidate assignments. |
| Statistical analyses | Physical-flowcell eligibility, contamination tests, regional model diagnostics, permutation results and all three breadth thresholds, plus reference-selection summaries and fixed-catalog-v2 host-depletion and capping comparisons. |
| Mycoplasmatales | Phylogenetic labels and supporting 16S comparisons. |
| Trematode comparisons | Direct trematode-marker comparisons: specimen and reference identities, alignment lengths, coverage, competing matches and execution provenance. |

**Table B.** Data supporting the reported analyses. Links lead to supporting data in the public analysis repository. The supporting-data provenance records the original analysis paths and checksums. The additional screening bundle reports initial screen nominations, their subsequent support and the SSU background-control comparison. See the Data Availability statement for the corresponding data and code deposits.

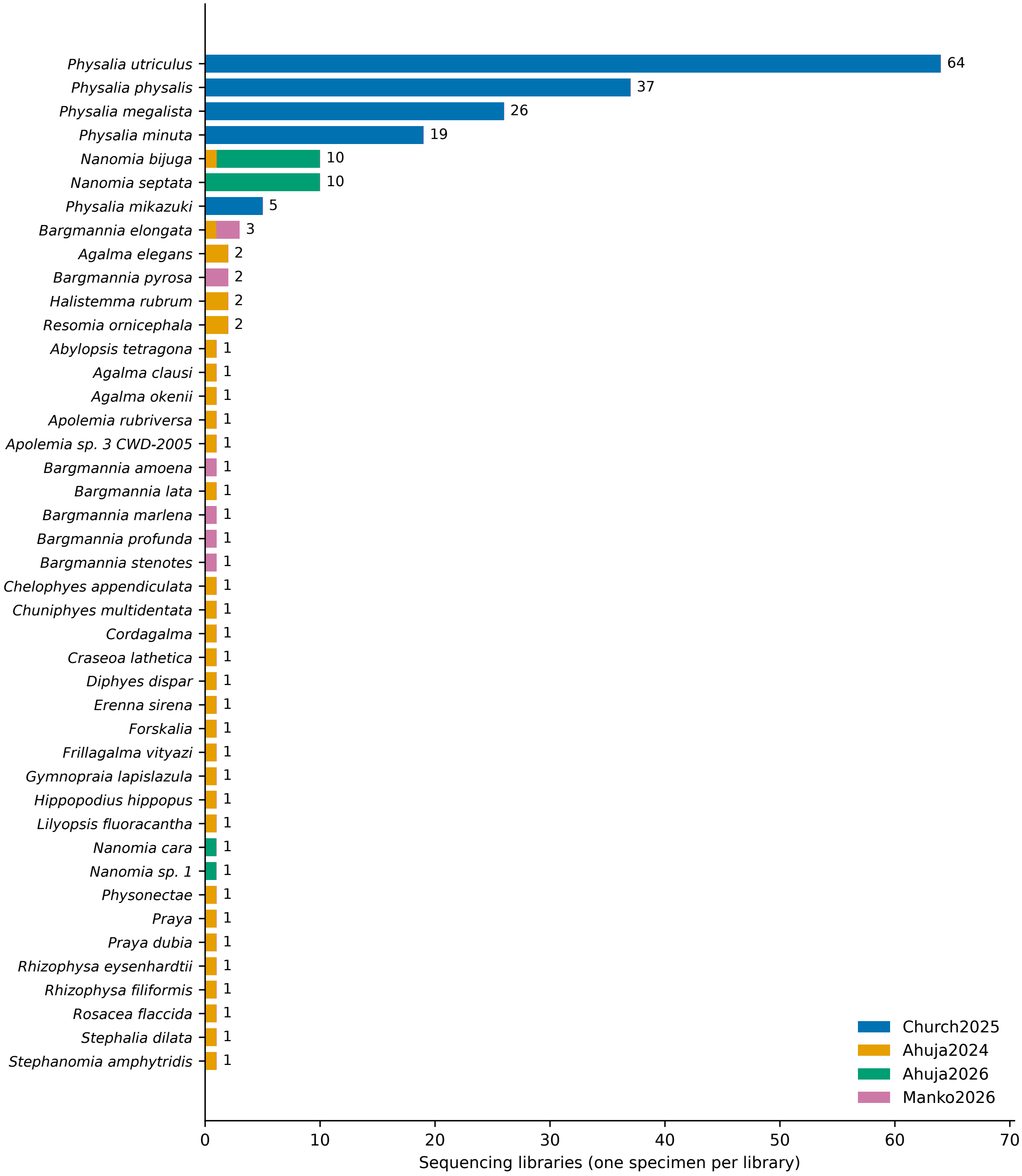


**Fig A.** Sampling was concentrated in *Physalia* and *Nanomia*. Bars show the number of specimens for each named taxon or unresolved identification across all 213 libraries, coloured by source study. The 43 bars represent 37 named taxa and 6 groups with broader or provisional identifications.

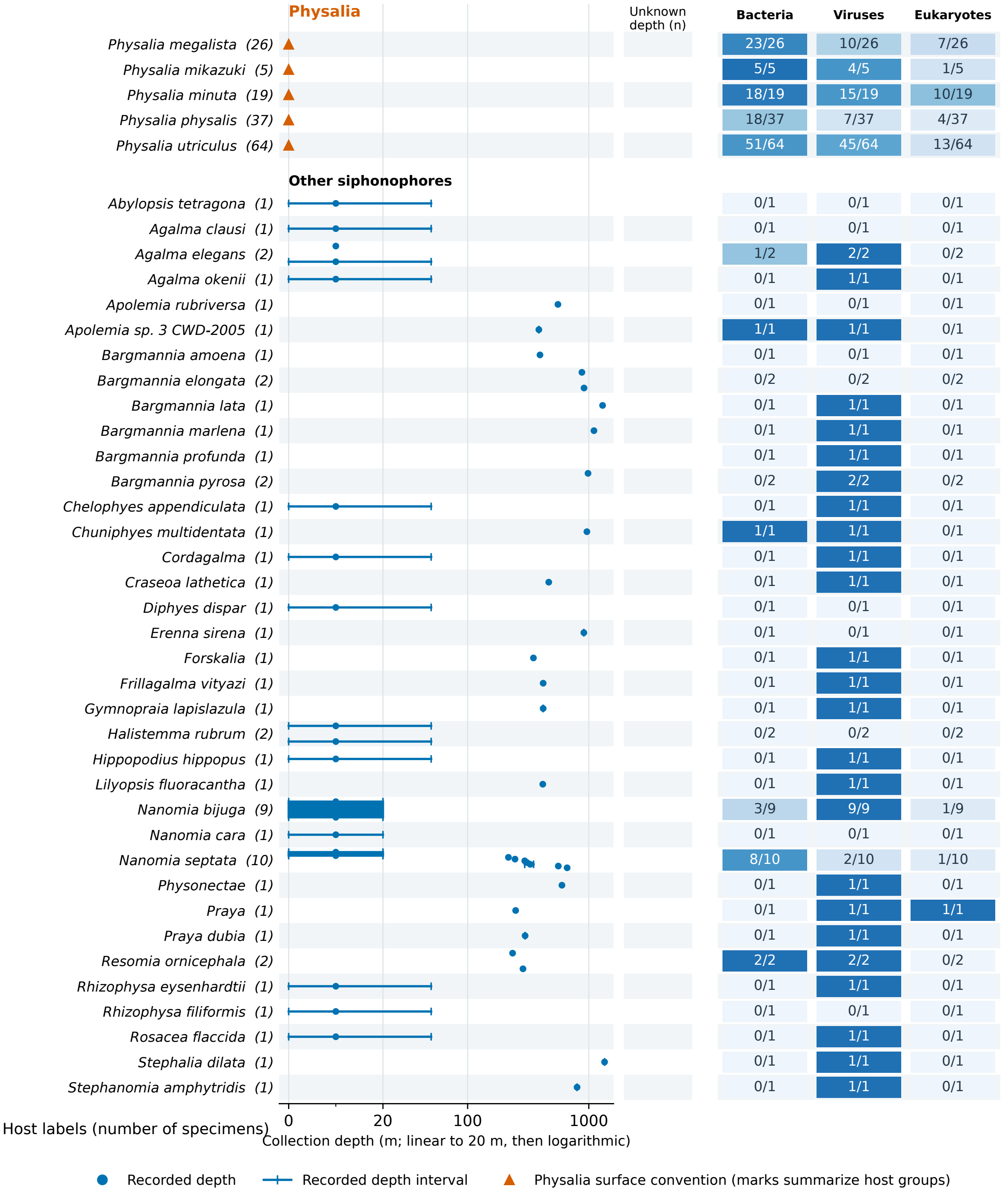


**Fig B.** Collection depths and detections across the 209 retained specimens. The surface-dwelling *Physalia* are summarized separately: one has a recorded 0 m depth and 150 have assigned surface values. For the other siphonophores, points and intervals retain recorded collection depths. Fractions give specimens with at least one accepted detection in each evidence category, with the number sampled as denominator. Host groups with broader or provisional identifications are retained. Frequencies do not represent equivalent measures of richness across bacteria, viruses and eukaryotes.

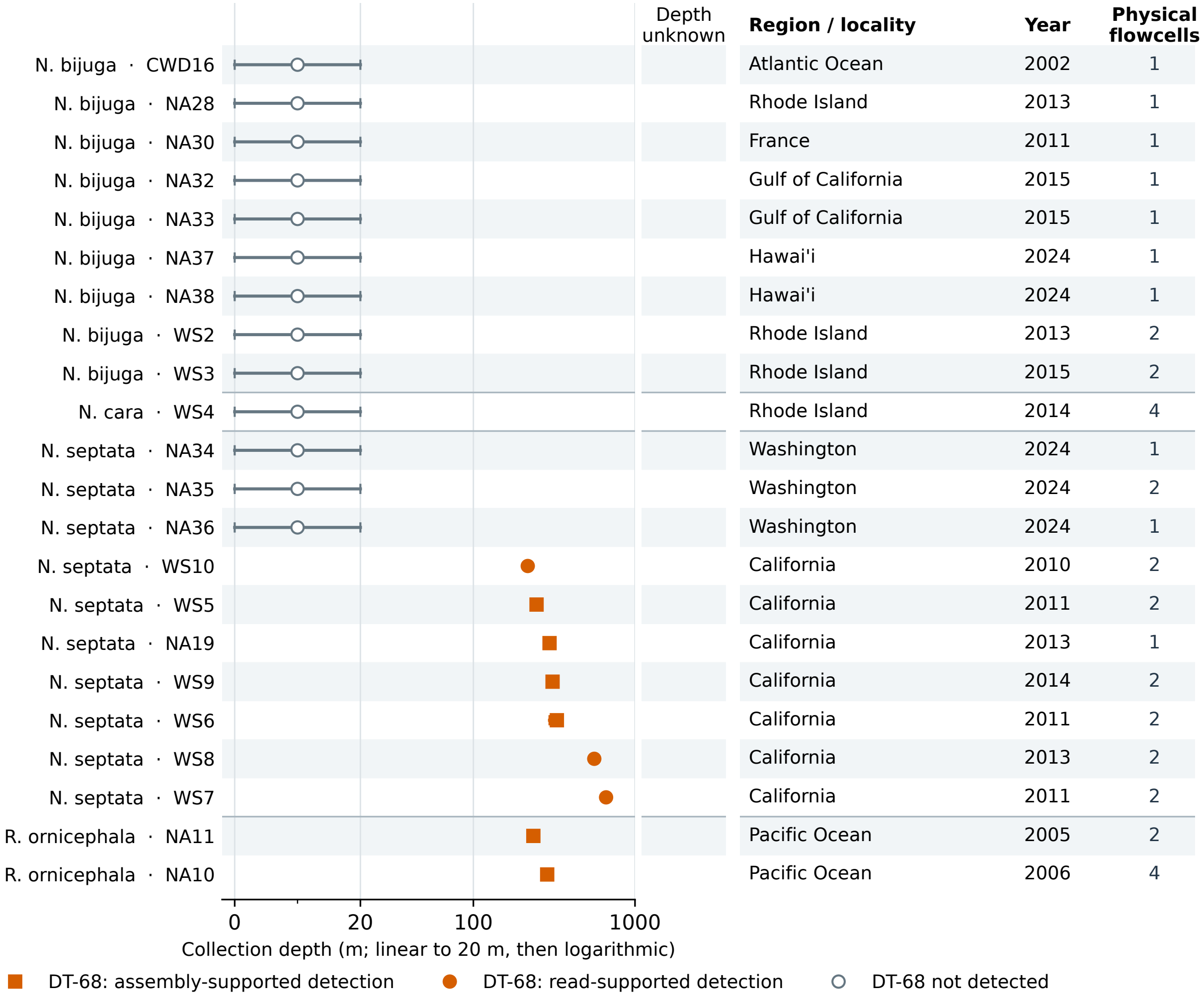


**Fig C.** A lineage of Metamycoplasmataceae bacteria, labelled DT-68 in the Genome Taxonomy Database, occurs in *Nanomia septata* and *Resomia* specimens collected at greater depths. The retained *Nanomia* and both *Resomia* specimens are shown. Symbols distinguish detections supported by reads, those also supported by same-specimen assemblies, and no accepted detection. Depth intervals retain their endpoints. Locality, collection year and physical-flowcell counts show that locality and depth are confounded within *N. septata*; these data do not isolate a depth effect.

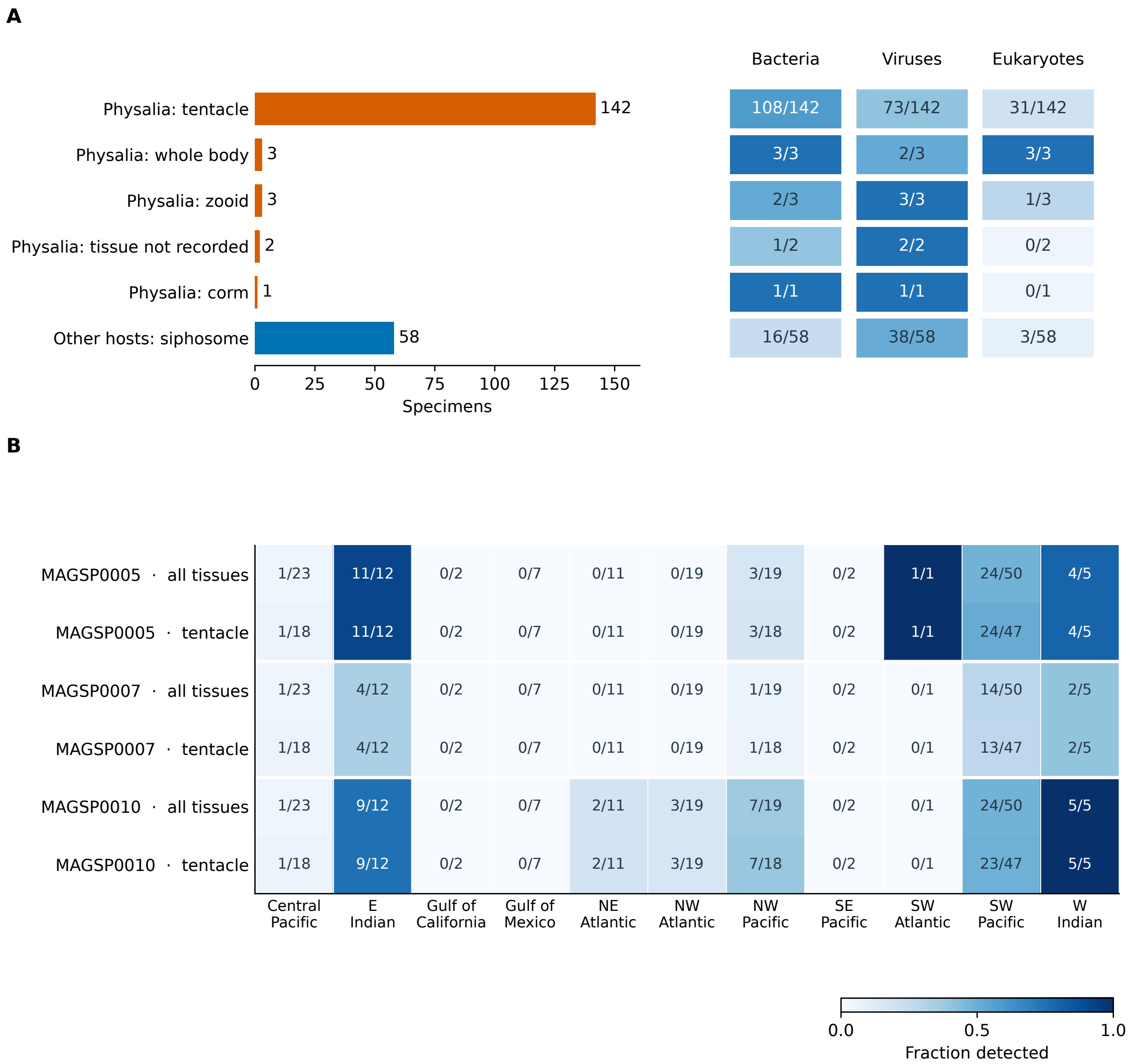


**Fig D.** Regional Metamycoplasmataceae differences persist within tentacle samples. (A) Recorded tissues for the 209 retained specimens, distinguishing *Physalia* tissues from siphosome samples in other siphonophores. Unknown tissues are labelled. (B) Detections of the three focal Metamycoplasmataceae genome clusters in all *Physalia* specimens and in recorded tentacle samples, with positive and total specimen counts. Regions without recorded tentacle samples have no estimate for that subset.

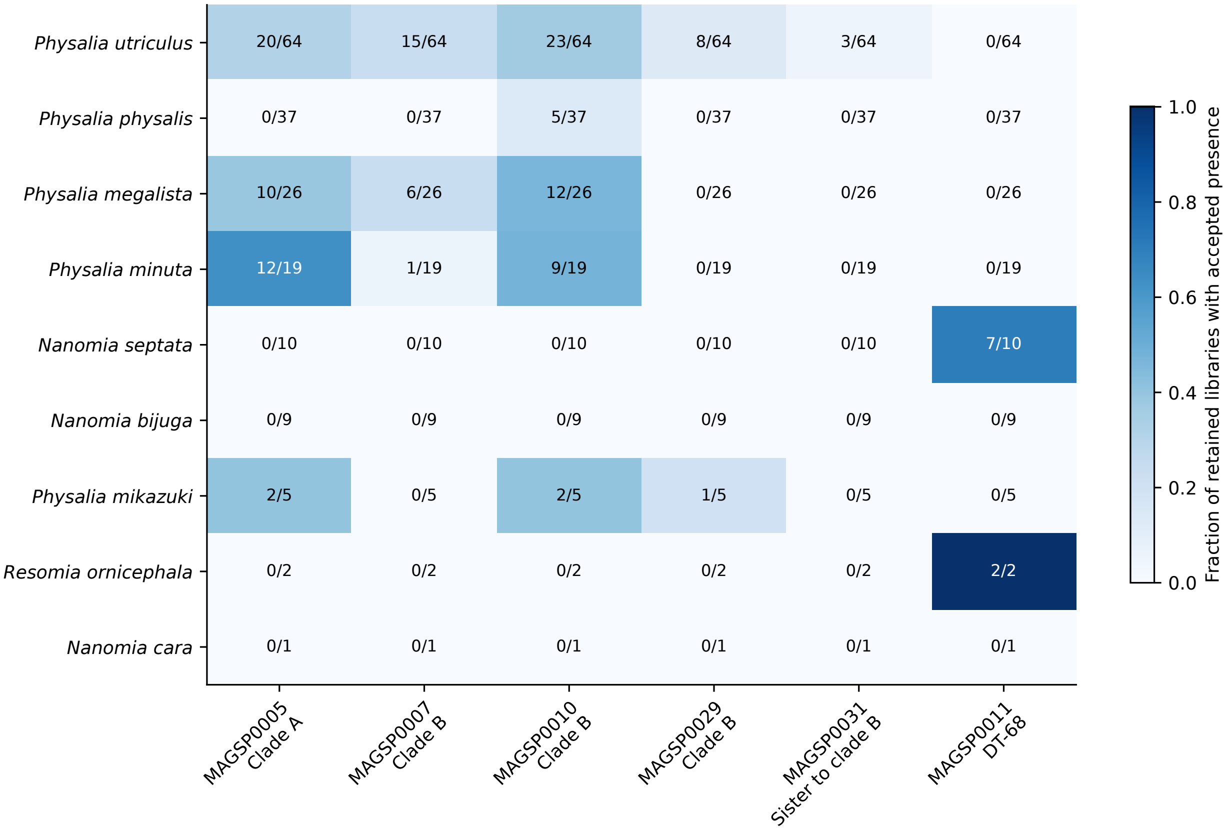


**Fig E.** Metamycoplasmataceae lineages differ in their siphonophore hosts. Each cell gives the number of libraries meeting the primary bacterial presence criterion over the total libraries for that host label; colour shows the corresponding fraction. All host groups with detections of these bacterial genome clusters are included.

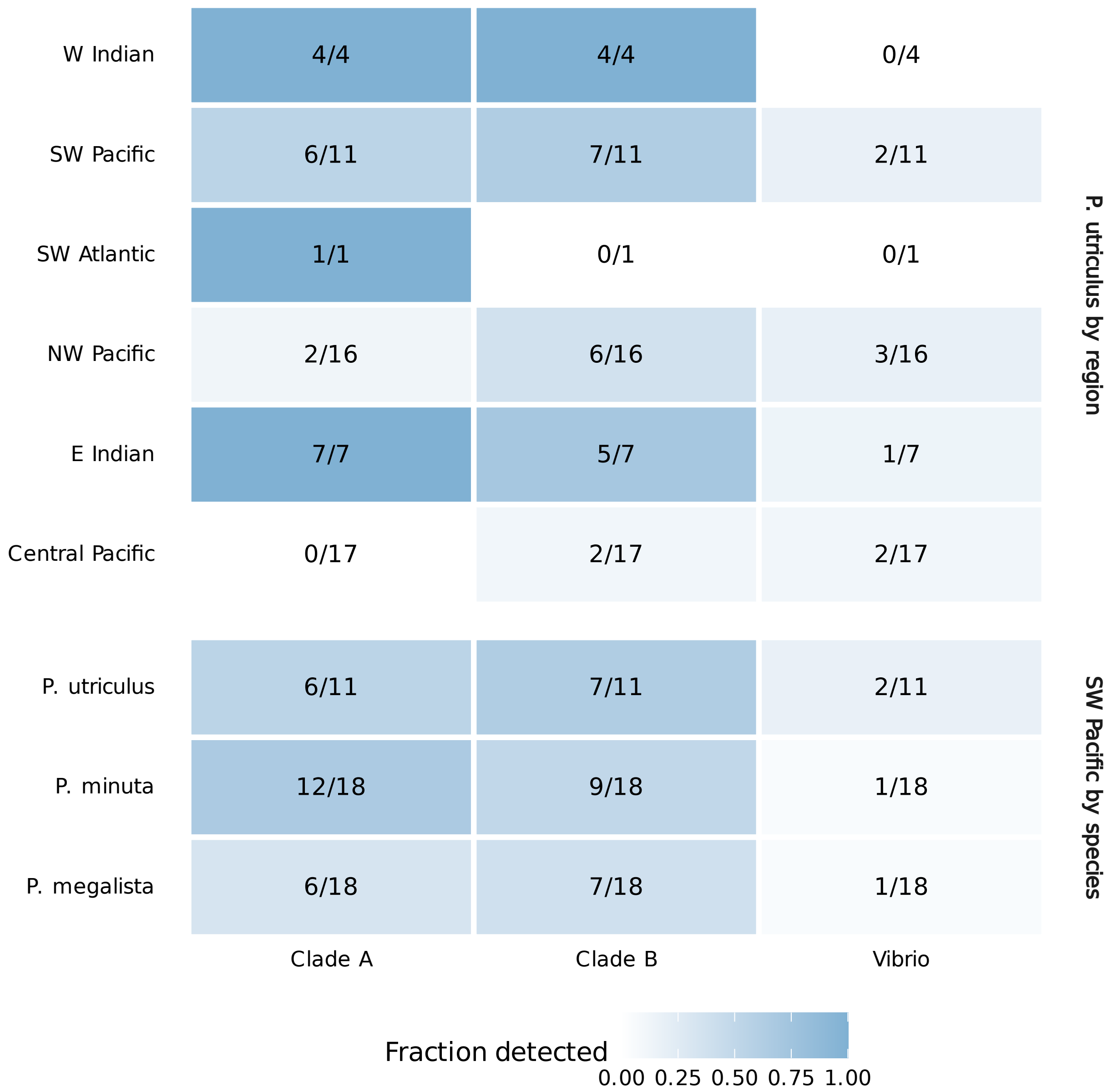


**Fig F.** Bacterial detections within *P. utriculus* across regions and among *Physalia* species within the southwestern Pacific, restricted to tentacle samples. Incidence includes all qualifying specimens; community tests use the single-flowcell subset with nonempty profiles. The two comparisons hold host identity or broad region constant, respectively.

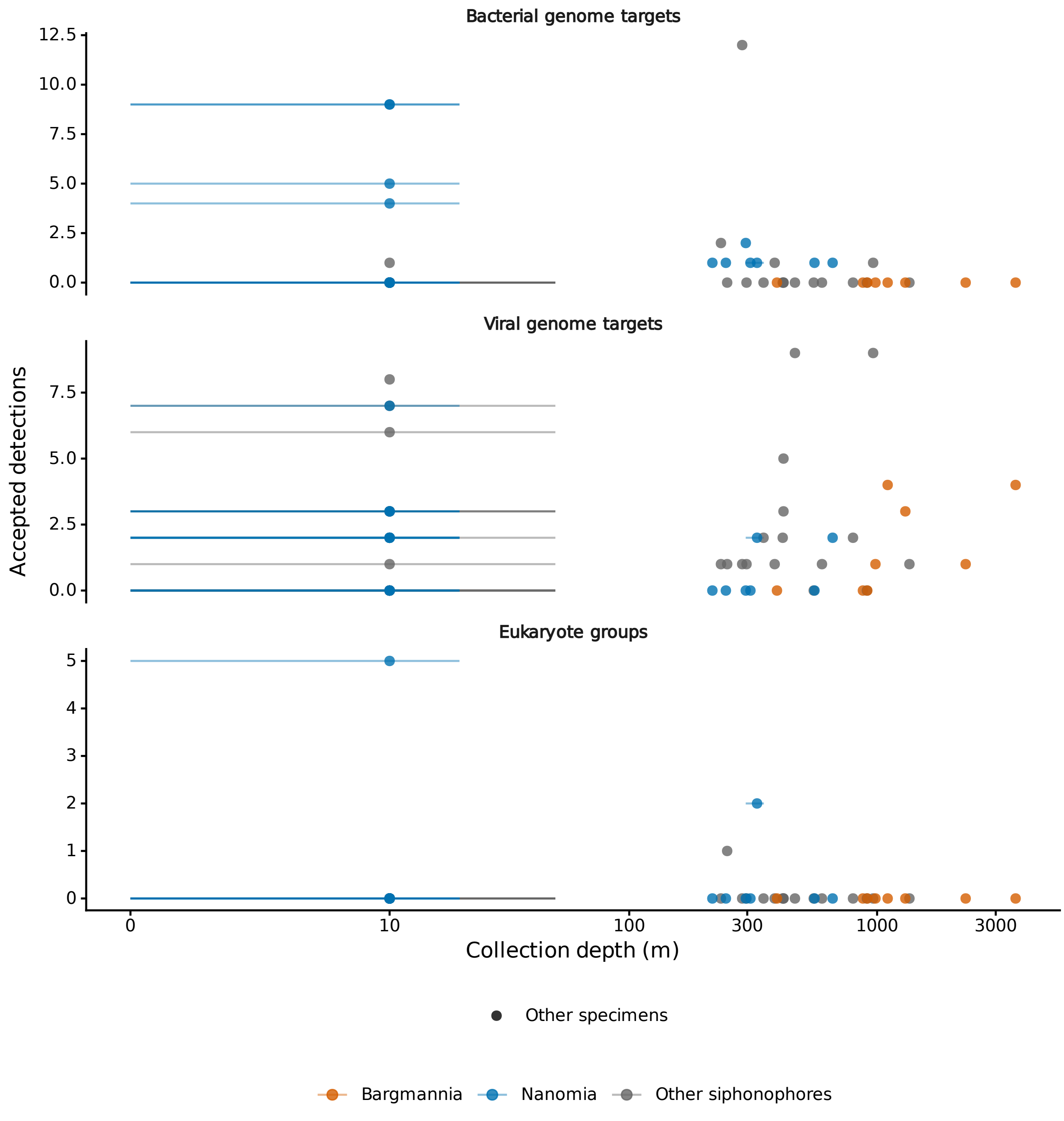


**Fig G.** Collection depth and associate detections in siphonophores other than *Physalia.* Intervals show recorded depth ranges. Detection counts use genome targets for microbes and broad reporting groups for eukaryotes. Colours distinguish *Nanomia*, *Bargmannia* and other siphonophores. Differences in host, locality and sampling constrain interpretation of depth effects.

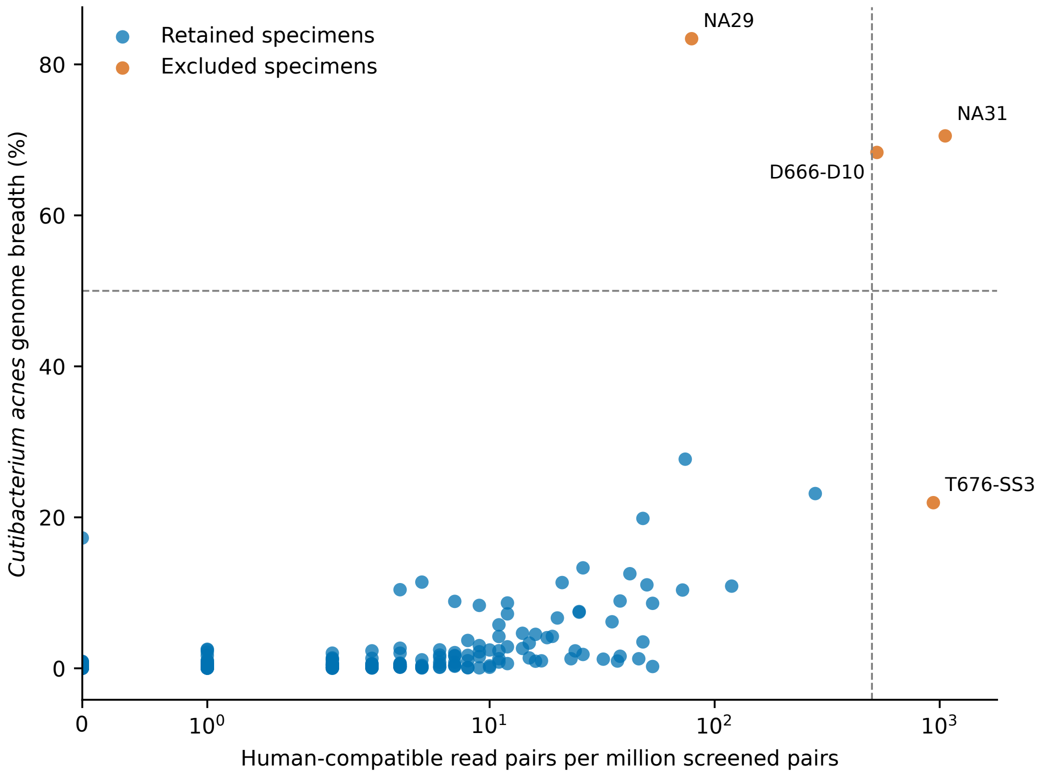


**Fig H.** Human-compatible reads and *Cutibacterium acnes* breadth identify libraries set aside from biological analyses. Each point is one of the 213 screened libraries. Axes show human-compatible read pairs per million screened pairs and the fraction of the *C. acnes* reference covered by reads. The four excluded libraries are labelled. Thresholds were chosen after inspecting gaps in the distributions: 500 human-compatible pairs per million (0.05%) or 50% *C. acnes* breadth. T676-SS3 also exceeded the SSU-control calibration. All *Physalia* libraries were retained.

### Author ORCIDs


| | |
|---|---|
| Casey W. Dunn | 0000-0003-0628-5150 |
| Namrata Ahuja | 0000-0002-2282-9755 |
| Dalila Destanović | 0000-0002-3744-5789 |
| Maciej K. Mańko | 0000-0001-6872-0256 |
| Martina Dal Bello | 0000-0003-3706-2929 |
| Samuel H. Church | 0000-0002-8451-103X |
